# A residual-ratio framework for auditing transcriptomic gene signatures against background expression structure

**DOI:** 10.64898/2026.04.11.717907

**Authors:** Ying Zhu, Chenyu Zhang, Vince D. Calhoun, Yuda Bi

## Abstract

**Background:** Transcriptomic gene signatures are widely used to infer pathway activity and biological mechanism from bulk cancer expression data, yet current evaluation strategies primarily emphasize internal coherence, predictive performance, or scoring robustness. A quantitative framework for assessing how much signature variation remains independent of background expression structure has been lacking.

**Results:** Unlike existing single-number signature-quality metrics such as Berglund uniqueness, residual-ratio auditing reports a *trajectory* across null-model richness: for each signature we compute the residual ratio 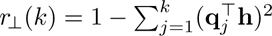 at progressively enriched expression-PC subspaces, together with an inverse-participation-ratio (IPR) concentration diagnostic that reports the effective number of axes absorbing each signature. Applied to a curated 17-entry benchmark, all 50 MSigDB Hallmark gene sets, and 1,181 Reactome pathways across 8 TCGA cancer types (4,462 samples), with external validation in METABRIC, the framework produces two complementary readouts. First, the curated panel is absorbed into the ExprPC50 subspace at residual ratios 18–43% below size-matched random 30-gene baselines in every cancer (curated mean *r*_⊥_ range 0.109–0.177 vs. random mean 0.182–0.288), providing the framework’s central quantitative discrimination between biologically coherent signatures and arbitrary gene combinations. Second, within the curated panel the ExprPC50 residual ratio is negatively correlated with the top-5 absorption concentration in every cancer (Spearman *ρ* from −0.59 in PRAD to −0.89 in SKCM, median −0.71; all 8 significant at *p <* 0.05, most at *p <* 10−3); we report this correlation as a descriptive geometric property of the null-model coordinate system rather than as a biological law, because 1,000 random 30-gene draws projected through the same top-50 expression-PC basis reproduce the same pan-cancer median *ρ* (−0.73; Supplementary Table S16), and it is robust to compositional nuisance: after rebuilding the null basis as immune-PC1 ⊕ stromal-PC1 ⊕ proliferation-PC1 plus 47 residual PCs, the per-cancer *ρ* becomes more negative rather than shallower (median −0.86; Supplementary Table S17), ruling out tumor purity, immune infiltrate, and stromal fraction as drivers of the pattern. Because absorption at ExprPC50 is a geometric property of how any signature direction sits in expression-PC space, tier-level distributional structure at this operating point is not separable beyond the low-vs-upper band split: a Kruskal–Wallis omnibus is significant (*p* = 4.9 × 10−13), but pairwise Dunn’s post-hoc tests show that Tiers 1, 4, and 5 are not separable (*p*_BH_ *>* 0.2). The trajectory shape itself is empirically *bootstrap-invariant*: across 200 sample-level fixed-basis bootstrap resamples of the 17 curated entries in BRCA, the mean pairwise Pearson correlation of trajectory-shape vectors is 0.999, and individual cell-level 95% bootstrap CI half-widths at *B*= 1,000 resamples are in the range 0.002–0.053. External replication in the METABRIC breast cancer cohort (*n*_samples_ = 1,980, microarray) showed moderate-to-strong rank-ordering concordance with TCGA-BRCA across the 17 curated entries (Spearman *ρ* = 0.72 on the 17-signature ordering, 95% Fisher-*z* CI 0.37–0.89, *p* = 0.001). Under an upper-bound sensitivity analysis, 45 of 50 Hallmark gene sets and 992 of 1,181 Reactome pathways had ExprPC200 residual ratios below the mean of their size-matched random baselines—a descriptive statistic reflecting axis alignment under rich null models, not a failure rate. In causal DAG simulations (*n*_rep_ = 100 replicates), a signature driven entirely by a latent confounder retained *r*_⊥_ = 0.233 at ExprPC50, numerically comparable to Tier 1 validated drivers, so a single-point residual ratio cannot adjudicate confounder-independence. The framework’s load-bearing signals are therefore the trajectory *shape* (statistically invariant under sample-level resampling) and the *magnitude gap* between the curated panel and its random 30-gene baseline (the curated-vs-random discrimination), read jointly—not the value of *r*_⊥_ at any single null-model dimensionality.

**Conclusions:** Residual-ratio auditing provides an interpretable and practical framework for quantifying how much of a transcriptomic gene signature’s variance remains orthogonal to a chosen background-expression model. The two statistically reliable quantities it reports are (i) the *shape* of the trajectory *r*_⊥_(*k*) across null-model richness, which is bootstrap-invariant across sample-level resamples, and (ii) the *magnitude gap* between the curated panel’s residual ratio and size-matched random 30-gene baselines at a fixed operating point, which is 18–43% in all 8 TCGA cancers and survives a purity-aware null-model construction. The negative correlation between *r*_⊥_ and the top-5 absorption concentration *c* (curated-panel median *ρ* = −0.71) is reproduced by random 30-gene sets under the same basis (random-draw median *ρ* = −0.73) and is therefore best read as a descriptive geometric property of the null-model coordinate system rather than a biological discovery about curated signatures. Any single operating-point residual ratio carries materially wider cell-level uncertainty than the trajectory shape and cannot, on its own, adjudicate confounder-independence. The framework’s outputs describe a signature’s geometric relationship to modeled background expression structure and do not evaluate clinical utility: a signature with a low residual ratio may still be clinically valuable when that low value reflects alignment with a strong prognostic or actionable program such as proliferation, immune infiltration, or cell cycle, and the framework is not a substitute for calibrated prognostic or predictive classifiers. All findings are based on bulk RNA-seq (TCGA PanCancer Atlas, 8 cancer types) and microarray (METABRIC) data; transfer to single-cell, single-nucleus, or spatial transcriptomics is out of scope and not claimed. Used within this scope—reading the trajectory shape and the magnitude-gap signal jointly, rather than the value of *r*_⊥_ at any one *k*—the framework adds a complementary audit layer to existing pathway-scoring and experimental-validation workflows, and supports more calibrated interpretation, comparison, and reporting of transcriptomic gene signatures in cancer studies.

## 1 Background

Transcriptomic gene signatures are one of the main interfaces through which cancer expression profiles are translated into pathway-level biological claims. They are routinely used to summarize pathway activity, tumor state, immune contexture, and therapy-relevant programs, and clinically mature examples such as the 70-gene prognostic profile, OncotypeDX, and PAM50 show that carefully developed signatures can be consequential for both biology and care [62–65]. Precisely because signatures sit at this interface between high-dimensional expression data and mechanistic interpretation, their apparent specificity matters.

In practice, signatures are often asked to do more than predict. They are used to support mechanistic narratives, stratify patients by putative pathway state, prioritize biomarkers, and interpret public cancer cohorts at scale. A signature can therefore appear useful while still primarily tracking a dominant transcriptomic axis, making interpretability a substantive issue for biological inference rather than a purely technical concern.

Several established tools evaluate adjacent aspects of signature quality. Cantini et al. classify signatures by internal coherence [77]; sigQC evaluates expression variability and scoring concordance [78]; and variancePartition decomposes variance at the gene level against user-specified covariates [82]. Closest to the present framework, Berglund et al. proposed a “uniqueness” metric for gene signatures based on the largest pairwise correlation between a signature score and any single global expression PC [83]; that metric returns one number per signature and does not vary the null-model basis. These methods address coherence, variability, scoring robustness, or single-PC redundancy, but none reports how much of a signature’s variance remains orthogonal to a *progressively enriched* background-expression subspace and how that orthogonal share evolves as the null model is enriched. The methodological norms for this kind of benchmarking have been articulated by Weber et al. [79], who emphasize null comparisons, fixed seeds, effect-size reporting, and public reproducible pipelines as non-negotiable ingredients of a credible methods benchmark; we adopt those norms throughout.

That missing layer matters because bulk transcriptomes are dominated by shared covariance structures. Proliferation, immune infiltration, stromal composition, purity, batch, and other global programs can explain large fractions of expression variance and can make pathway-level signatures appear more specific than they are if the signature largely recapitulates those axes. A framework intended to support mechanistic interpretation therefore needs to distinguish signal that is merely aligned with dominant transcriptomic structure from signal that remains distinguishable after that structure is modeled.

The methodological landscape for adjusting bulk expression for compositional structure is closely related but operationally distinct. Tumor purity and stromal fraction estimators such as ESTIMATE [22] and ABSOLUTE [88], and cell-type deconvolution tools such as CIBERSORT [23], xCell [24], and quanTIseq [89], estimate compositional fractions from bulk expression that can then be used to adjust downstream analyses. Residual-ratio auditing asks a complementary question: regardless of which biological axes populate the null-model subspace, how much of a signature’s variance remains orthogonal to that subspace? Deconvolution estimates cell-type fractions and is maximally informative when the goal is to decompose bulk expression into its compositional contributors; residual-ratio auditing quantifies how much of a signature’s variance is distinguishable from the dominant axes that those contributors populate, and is therefore a natural second-layer check for studies that make pathway-level interpretation claims from pre- or post-deconvolution expression.

Prior work already points at pieces of this problem: random gene sets can predict outcome in breast cancer [6], prognostic markers are often enriched for housekeeping and cell-cycle genes [7, 81], and batch effects, purity, and dimensionality reduction can reshape inferred biological structure [9, 15, 17]. Taken together, these observations do not imply that transcriptomic signatures are uninformative. They imply that the field needs a quantitative way to ask, for a given signature in a given dataset, how much of its variance remains orthogonal to a specified background-expression subspace—and how that orthogonal share changes as the subspace is enriched from a single interpretable axis (proliferation PC1) to the leading principal components of the full expression matrix.

Here we introduce a residual-ratio framework to measure that quantity directly. In plain terms, the *residual ratio* asks what fraction of a signature’s sample-to-sample variation remains after projection away from a chosen background-expression model, whereas *absorption concentration* asks whether the absorbed variance is dominated by one axis or distributed across many. This separates three questions that are often conflated in practice: whether a signature is active, whether it is clinically useful, and whether its variation is distinguishable from modeled background transcriptomic covariance. At panel scale, we will show that these two quantities have a distinctive joint structure: the curated panel is absorbed into the ExprPC50 subspace at residual ratios 18–43% below size-matched random 30-gene baselines in every cancer, which is the framework’s central quantitative finding, and—as a separate descriptive observation—the negative correlation between the ExprPC50 residual ratio and the top-5 absorption concentration (median Spearman *ρ* = −0.71) is a property of how *any* gene-set direction aligns with the leading expression-PC axes. Random 30-gene draws reproduce the same pan-cancer median (Supplementary Table S16), so this correlation is reported as a coordinate-system consistency check rather than as a biological-validation property.

We benchmark this framework across 16 curated pathway signatures, one housekeeping control, all 50 MSigDB Hallmark gene sets, and 1,181 Reactome pathways in 8 TCGA cancer types, with external validation in the METABRIC breast cancer cohort and data access through PanCancer Atlas resources such as cBioPortal [1–5]. We use ExprPC50 as the primary interpretive operating point—a design choice supported by cross-validated dimensionality selection (the strict CV optimum in BRCA and inside the CV-admissible ranking-stability range in 6 of the 7 other primary cancers; Methods §Cross-validated dimensionality selection)—and report ExprPC200 as an upper-bound sensitivity analysis, then examine representative exemplars, curated-panel structure, public-pathway scale, gene-overlap effects, and causal simulations to define where residual-ratio auditing is most informative and where its interpretation should remain bounded. This framework is intended to complement pathway scoring and experimental validation by adding a practical audit layer that makes a signature’s relationship to dominant background expression axes explicit and reportable.

## 2 Results

Unless noted otherwise, primary interpretation refers to ExprPC50, and ExprPC200 is treated as an upper-bound sensitivity analysis.

### 2.1 Framework overview and benchmark design

Residual-ratio auditing measures how much signature variance remains outside progressively richer models of background expression structure. Throughout, we use *absorbed* as the complementary view of *residual*: for any signature direction **h** and null-model subspace *𝒯*, the absorbed fraction equals 1 − *r*_⊥_ and represents the share of **h** that lies inside 𝒯; a high residual ratio is therefore equivalent to low absorption, and vice versa. To make this quantity interpretable rather than purely abstract, we audited each signature against a hierarchy of null-model subspaces 𝒯 ranging from a single proliferation principal component (dim(𝒯) = 2) to the top 200 principal components of the full expression matrix (dim(𝒯) = 201). Lower levels of the hierarchy ask whether a signature is already explained by dominant biological axes such as proliferation or immune infiltration, whereas richer expression-PC nulls ask how quickly the same signature is absorbed by global transcriptomic covariance. The top 10 expression PCs are not “agnostic” axes: in every cancer at least one of PC1–PC3 tracks an immune, stromal, or proliferation proxy at |*ρ*| ≥ 0.5, and the leading PC1 is dominated by a single compositional program in 5 of 8 cohorts (e.g., COAD PC1 is essentially stromal at *ρ* = −0.86, OV PC1 is immune at −0.72, SKCM PC1 is immune at +0.70; full 8×10×3 annotation in Supplementary Table S19). This biological content is the source of axis interpretability in the hierarchy and is *not* a confound: Supplementary Table S17 shows that explicitly projecting out an immune ⊕ stromal ⊕ proliferation anchor block and rebuilding the null on the residual expression matrix leaves the pan-cancer *ρ*(*r*_⊥_*, c*^(5)^) correlation intact (in fact deepening from −0.712 to −0.857).

We applied this hierarchy to a benchmark comprising 16 curated pathway signatures, one housekeeping control, all 50 MSigDB Hallmark gene sets, and 1,181 Reactome pathways across 8 TCGA cancer types (4,462 samples) (Figure 1, Table 1). For each signature, we computed the residual ratio *r*_⊥_ = ‖**R**‖2*/*‖**h**‖2, complemented when useful by absorption concentration, and compared the observed values against matched random baselines. This benchmark was designed to support both method validation and practical interpretation: curated pathway signatures provide biologically recognizable use cases, the housekeeping control anchors a non-pathway reference, public pathway collections test generality, and the hierarchy itself reveals whether absorption occurs early along interpretable axes or only under richer agnostic null models. The cross-cancer heatmap in Figure 1c already shows that the benchmark is not organized by a single smooth continuum: high-residual and low-residual regimes recur, but their relative spacing depends on cancer context. The next question is therefore whether this structure is stable enough to support a primary operating point for interpretation.

**Figure 1:**
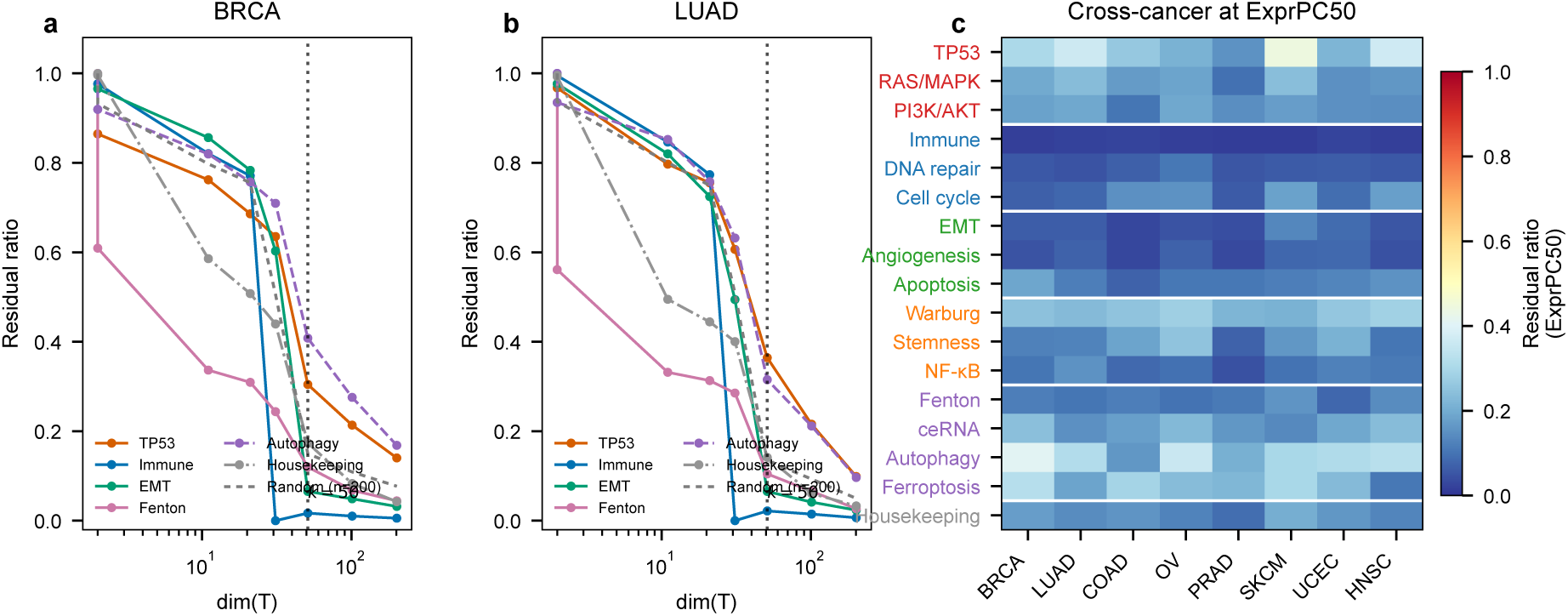
Framework overview and benchmark design. (**a**) Residual-ratio trajectories for representative signatures in BRCA across the null-model hierarchy. The dashed gray line shows the mean *r*_⊥_ of 200 random 100-gene sets and the vertical dotted line marks the primary interpretive operating point (*k* = 50). (**b**) Residual-ratio trajectories for the same representative signatures in LUAD. (**c**) Cross-cancer heatmap of *r*_⊥_ at ExprPC50 for the 16 curated pathway signatures and the housekeeping control across 8 TCGA cancer types. Tier separators (white lines) group the pathway signatures by validation tier.

**Table 1:** Residual ratios at the primary ExprPC50 operating point across 8 TCGA cancer types. Bold values indicate the highest residual ratio per cancer type. The “Random” row shows the mean ratio of 200 random 100-gene sets. ExprPC50 is the primary interpretive operating point (Methods §Cross-validated dimensionality selection; design choice rather than strict CV optimum). Sample-only bootstrap 95% confidence intervals for the BRCA and LUAD columns (1,000 fixed-gene-space-basis sample-level resamples with replacement; see Methods § Statistical analysis) are reported in Supplementary Table S5: the 1,000-replicate procedure yields 95% CI half-widths in the range ∼0.002–0.053 (BRCA median ≈ 0.019 LUAD median 0.024) tighter than the earlier 200-replicate report by roughly 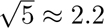 and are the numbers to read when comparing cell-level contrasts. The corresponding upper-bound ExprPC200 summary is reported in Supplementary Table S3.

| Tier | Signature | BRCA | LUAD | COAD | OV | PRAD | SKCM | UCEC | HNSC |
| --- | --- | --- | --- | --- | --- | --- | --- | --- | --- |
| 1 | TP53 pathway | 0.304 | **0.364** | 0.262 | 0.214 | 0.154 | **0.442** | 0.220 | **0.368** |
| 1 | RAS/MAPK | 0.198 | 0.235 | 0.171 | 0.185 | 0.097 | 0.240 | 0.151 | 0.163 |
| 1 | PI3K/AKT/mTOR | 0.183 | 0.192 | 0.104 | 0.193 | 0.148 | 0.173 | 0.151 | 0.147 |
| 2 | Immune checkpoint | 0.017 | 0.022 | 0.024 | 0.018 | 0.015 | 0.013 | 0.024 | 0.017 |
| 2 | BRCA DNA repair | 0.058 | 0.047 | 0.067 | 0.106 | 0.053 | 0.059 | 0.063 | 0.058 |
| 2 | Cell cycle/CDK | 0.067 | 0.081 | 0.151 | 0.152 | 0.062 | 0.184 | 0.092 | 0.180 |
| 3 | EMT | 0.065 | 0.065 | 0.028 | 0.049 | 0.041 | 0.135 | 0.089 | 0.054 |
| 3 | Angiogenesis | 0.047 | 0.070 | 0.030 | 0.070 | 0.035 | 0.079 | 0.083 | 0.045 |
| 3 | Apoptosis/BCL2 | 0.188 | 0.119 | 0.073 | 0.108 | 0.106 | 0.118 | 0.132 | 0.153 |
| 4 | Warburg effect | 0.249 | 0.233 | 0.254 | 0.278 | 0.220 | 0.211 | 0.258 | 0.282 |
| 4 | Stemness | 0.122 | 0.125 | 0.186 | 0.251 | 0.072 | 0.160 | 0.211 | 0.104 |
| 4 | NF-κB inflammatory | 0.103 | 0.154 | 0.083 | 0.094 | 0.045 | 0.099 | 0.122 | 0.110 |
| 5 | Fenton metabolic | 0.120 | 0.105 | 0.120 | 0.100 | 0.112 | 0.157 | 0.081 | 0.144 |
| 5 | ceRNA hubs | 0.245 | 0.150 | 0.176 | 0.229 | 0.160 | 0.138 | 0.194 | 0.237 |
| 5 | Autophagy | **0.408** | 0.315 | 0.162 | **0.359** | 0.211 | 0.299 | **0.310** | 0.324 |
| 5 | Ferroptosis | 0.322 | 0.186 | **0.292** | 0.235 | **0.223** | 0.298 | 0.231 | 0.107 |
| C | Housekeeping | 0.174 | 0.141 | 0.157 | 0.128 | 0.092 | 0.205 | 0.161 | 0.132 |
| C | Random ($n = 200$) | 0.150 | 0.134 | 0.088 | 0.187 | 0.080 | 0.110 | 0.122 | 0.095 |

### 2.2 Validation and robustness identify ExprPC50 as a practical operating point

The ranking structure remained stable across the main analytic choices that typically perturb gene-signature studies, supporting ExprPC50 as a practical primary operating point rather than an arbitrary convenience (Figure 2). Across 12 alternative null constructions spanning biological covariates and agnostic expression-PC families, signature rankings were highly stable within the expression-PC family (median pairwise Spearman *ρ* = 0.983 in BRCA and *ρ* = 0.953 in LUAD). Four scoring methods—uniform weight 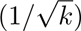, z-score, mean expression, and rank-based—also produced highly concordant rankings, with a minimum pairwise Spearman *ρ* = 0.912 and all other pairs above *ρ* = 0.95.

**Figure 2:**
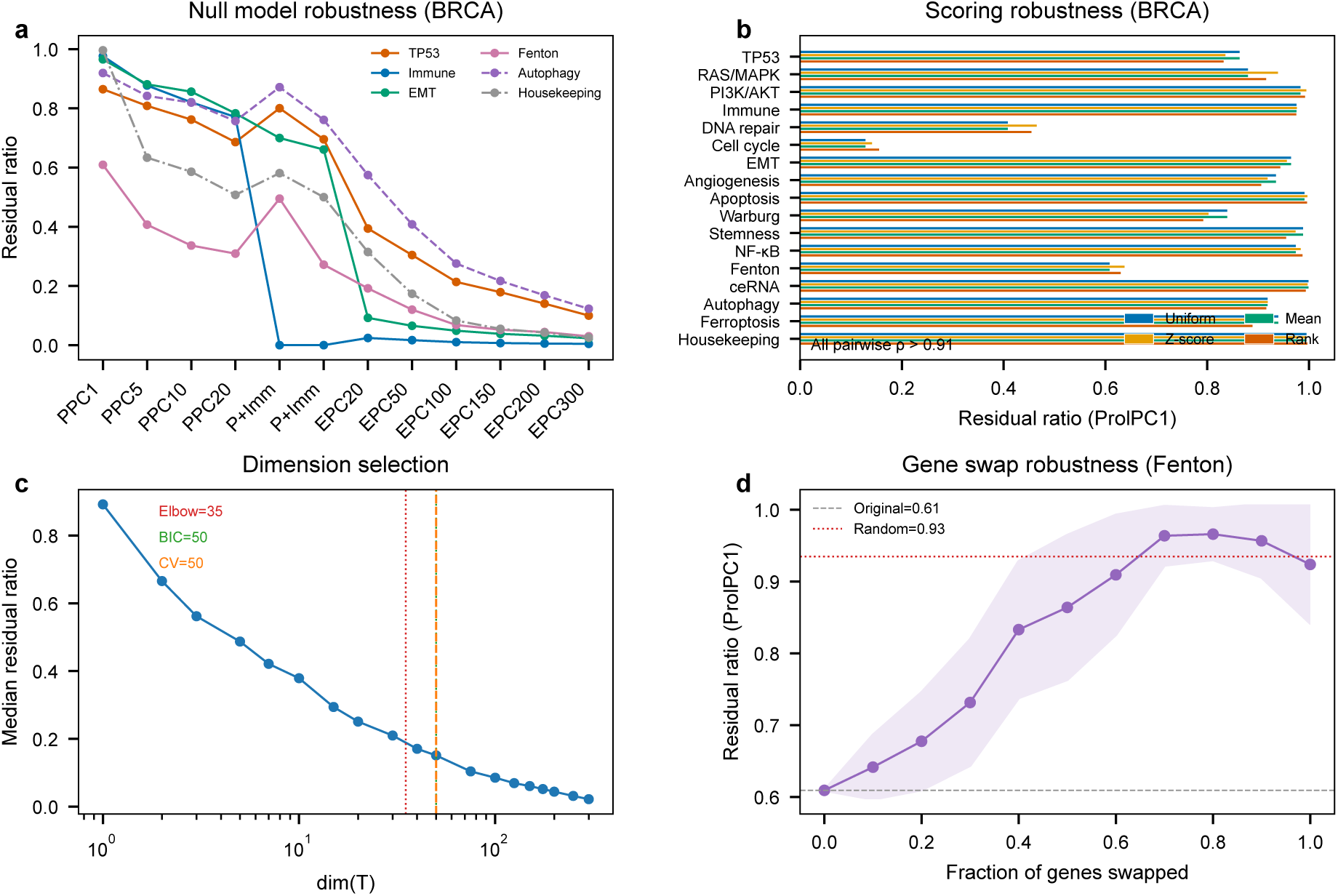
Validation and robustness of the auditing framework. (**a**) Null-model robustness for representative signatures in BRCA across 12 alternative null constructions spanning biological covariates and expression-PC families. (**b**) Scoring robustness in BRCA across four signature-scoring schemes; the panel annotation summarizes pairwise rank-correlation concordance. (**c**) Cross-validated dimensionality selection showing the consensus operating point at *k* = 50 together with elbow and BIC heuristics. (**d**) Gene-swap robustness for the Fenton metabolic signature in BRCA with the observed ProlPC1 residual ratio compared against the trajectory obtained as signature genes are progressively replaced.

Dimensionality selection supports ExprPC50 as a natural operating point in BRCA and as a reasonable choice across the other 7 cancers. A 5-fold cross-validation sweep across *k* ∈ {5, 10, 20, 50, 100, 200, 300} with an out-of-sample gene-space projection and per-fold QR re-orthonormalization selected *k*^∗^ = 50 as the strict CV optimum in BRCA (train–test gap 0.025, ranking stability *ρ* = 0.98; Supplementary Table S12, Methods §Cross-validated dimensionality selection), and *k*^∗^ = 20 in LUAD, COAD, PRAD, UCEC, and HNSC; SKCM prefers *k*^∗^ = 5 and OV prefers *k*^∗^ = 200. ExprPC50 lies inside the CV-admissible range (ranking stability *ρ* ≥ 0.87) for seven of the eight primary-analysis cancers, and is the strict CV optimum for BRCA, the cohort on which the main exemplar analysis is anchored. We therefore use ExprPC50 as the primary interpretive operating point across all cancers and report ExprPC200 as an upper-bound sensitivity level. The same qualitative ordering survives size-, mean-expression-, and variance-matched random baselines, indicating that the principal conclusions are not artifacts of a single baseline choice.

External validation and perturbation tests supported both breast-cancer cross-cohort concordance and signature specificity of the ranking. In the independent METABRIC breast cancer cohort (*n*_samples_ = 1,980, microarray), the ordering of the 16 curated pathway signatures together with the housekeeping control showed moderate-to-strong concordance with TCGA-BRCA (Spearman *ρ* = 0.72 across the *n* = 17 curated benchmark entries—the unit of analysis is the 17-signature ordering rather than per-sample data, so the Fisher-*z* CI and permutation *p*-value below are computed on the 17-pair rank correlation, not on *n* = 1,980 samples—95% Fisher-*z* CI 0.37–0.89, *p* = 0.001; the lower CI bound admits a moderate rather than a very-strong effect at this benchmark size); the upper-band members (TP53, *r*_⊥_ = 0.43; PI3K/AKT/mTOR, 0.38; autophagy, 0.33) again occupied the top of the benchmark at ExprPC50, whereas the immune checkpoint signature again showed the lowest residual ratio (*r*_⊥_ = 0.02). Additional gene-swap analyses showed that the low residual ratios of metabolism-associated signatures were driven by gene identities rather than trivial effects of gene-set size, mean expression, or variance, and within-subtype BRCA analyses indicated that a substantial share of absorption is carried by between-subtype structure rather than residual within-subtype noise. With the operating point established, the next question is what distinct absorption regimes actually look like in representative signatures.

### 2.3 Worked examples reveal distinct absorption regimes

Three representative signatures illustrate three recurring absorption patterns when traced across the null-model hierarchy (Figure 3b,c; Table 1). At the primary ExprPC50 operating point, TP53 sits in the upper residual-ratio band (cross-cancer median *r*_⊥_ = 0.283, range 0.154–0.442); the immune checkpoint signature defines a low-residual few-axes regime (median 0.018, range 0.013–0.024); and the Fenton metabolic signature illustrates early proliferation-axis absorption at ProlPC1, where it drops to *r*_⊥_ = 0.61 in BRCA, although by ExprPC50 its residual ratio (median 0.116) sits close to the random baseline rather than at the most absorbed extreme. These three patterns characterize distinct *trajectories* of variance across null-model levels rather than ranking the signatures themselves, and we use them below as a practical vocabulary for reading the full benchmark.

**Figure 3:**
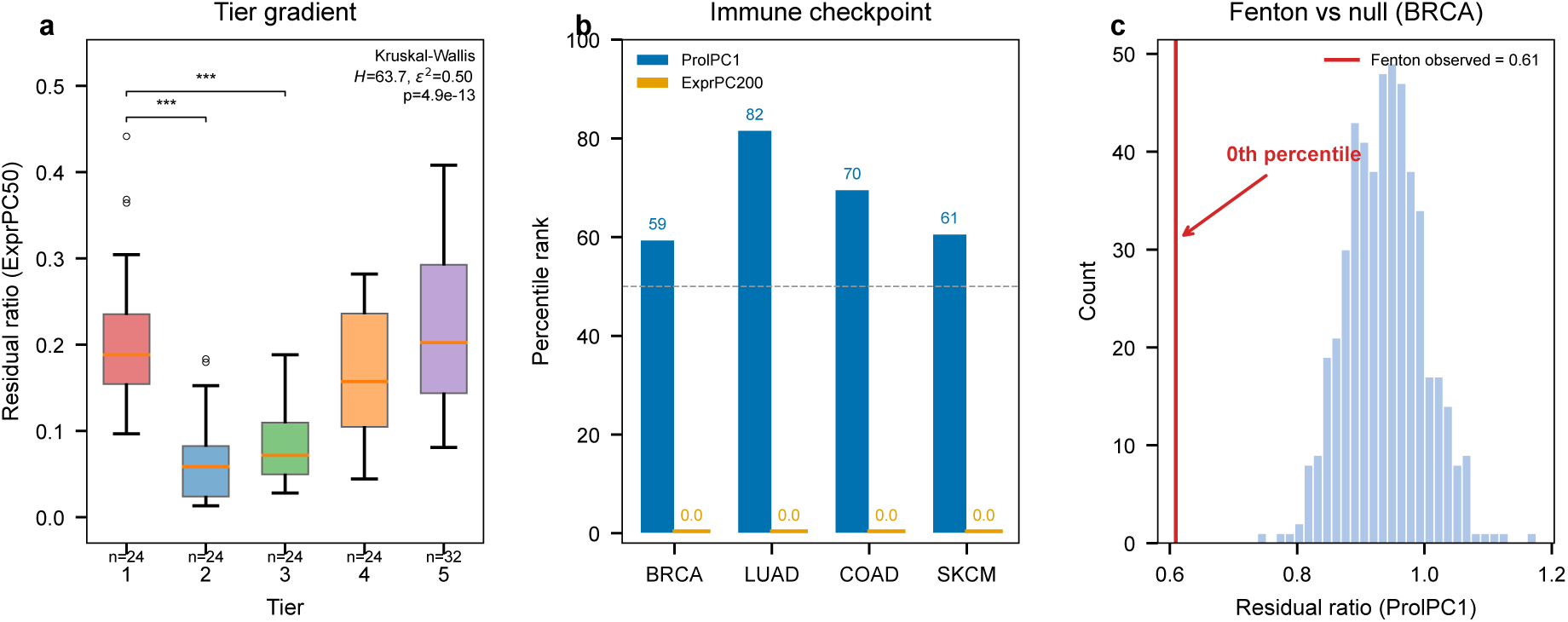
Worked examples and curated-panel synthesis. (**a**) Tier gradient at ExprPC50 aggregated across all 8 cancer types. Box plots show residual ratios for the 16 curated pathway signatures grouped by validation tier (Tier 1–4: *n* = 24 each; Tier 5: *n* = 32). Panel annotation reports the across-tier Kruskal–Wallis test (*H* = 63.7, *ε*^2^ = 0.50, *p* = 4.9 10−13); pairwise Dunn’s post-hoc BH-corrected significance is reported in Supplementary Table S4. (**b**) Immune checkpoint percentile ranks versus size-matched random controls in four cancer types, contrasting the proliferation-only ProlPC1 null with the upper-bound ExprPC200 sensitivity level. Exact numeric values are printed next to each bar because the ExprPC200 percentile ranks are uniformly near zero—a scale mismatch with the ProlPC1 ranks that is itself the panel’s intended message. (**c**) Fenton metabolic residual ratio in BRCA relative to its ProlPC1 null distribution, showing early absorption and a 0th-percentile ranking versus matched random controls.

TP53 illustrates a persistent-orthogonal regime in which substantial variance remains outside the modeled background structure across the full null-model hierarchy. Even at the weakest BRCA null model (proliferation PC1 only), TP53 retains 87% of its variance (*r*_⊥_ = 0.87), and at ExprPC50 it occupies the upper residual band alongside both validated drivers such as RAS/MAPK and PI3K/AKT and independently validated cellular programs such as autophagy and ferroptosis (Table 1). This is the pattern expected when a signature captures variation that is not reducible to a single dominant background axis.

The immune checkpoint signature illustrates a different regime: it is not proliferation-driven, but it is highly sensitive to immune-aware and richer expression-PC nulls. At ProlPC1 it ranks between the 59th and 85th percentile versus size-matched random controls across BRCA, LUAD, COAD, and OV, showing that proliferation alone does not explain it. Under immune-enriched or richer expression-PC nulls, however, the same signature becomes one of the most strongly absorbed entries in the benchmark, indicating that its signal is concentrated along a small number of dominant immune-related axes rather than distributed broadly outside the null space.

Fenton metabolic provides the clearest example of *early proliferation-axis* absorption in this benchmark. At the weakest BRCA null model (ProlPC1), it already falls to *r*_⊥_ = 0.61, and it reaches the 0th percentile versus size-matched random controls in BRCA, LUAD, COAD, and OV at ProlPC1. In this setting, most of the signature’s variance is captured almost immediately by a proliferation-associated background axis. At the primary ExprPC50 operating point the Fenton residual ratio (cross-cancer median 0.116) sits close to the random-baseline mean (0.116), because nearly all of the variance that a proliferation-only null can capture has already been captured by ProlPC1 itself and little additional absorption occurs as the null model is enriched from ProlPC1 to ExprPC50. Together, these three examples provide a practical vocabulary for reading the broader benchmark: distributed high residual (TP53), few-axis low residual (immune checkpoint), and early proliferation-axis absorption (Fenton at ProlPC1).

### 2.4 Curated-panel synthesis places the exemplars in context

The curated panel’s primary quantitative signal is a consistent absolute gap in residual ratio relative to size-matched random baselines. Across 1,000 random 30-gene draws per cancer, the 17-entry curated panel’s mean ExprPC50 residual ratio is between 0.109 (PRAD) and 0.177 (SKCM), while the random-draw mean is between 0.182 (PRAD) and 0.288 (OV); the curated panel is therefore absorbed into the ExprPC50 subspace at residual ratios 18–43% below the random baseline in every cancer (Supplementary Table S16). This absolute-gap finding is the framework’s central discrimination between biologically coherent signatures and arbitrary gene combinations, and is what separates the curated panel from the null.

A second, descriptive observation about the curated panel is that, within it, the ExprPC50 residual ratio is negatively correlated with the top-5 absorption concentration statistic in every cancer (Supplementary Table S10, Methods § 4.7): the Spearman rank correlation ranges from *ρ* = −0.59 in PRAD (*p* = 1.3 × 10−2) to *ρ* = −0.89 in SKCM (*p* = 1.7 × 10−6), with a median of −0.71 across cancers; the corresponding correlations with the parameter-free inverse-participation ratio (IPR) are positive in every cancer, range from *ρ* = +0.51 in BRCA to +0.86 in SKCM, median +0.62, and all 8 pan-cancer correlations are significant at *p <* 0.05 (six at *p <* 10−3). This correlation is not confounded by gene-set size: computing the partial Spearman *ρ*(*r*_⊥_*, c*^(5)^ | *n*_genes_) via rank-residual regression on the 17 curated entries (whose gene counts span 28–66, median 30) leaves the per-cancer correlation essentially unchanged or slightly more negative (median shift −0.022; Supplementary Table S18).

We report this correlation as a descriptive geometric property of the null-model coordinate system rather than as a biological discovery about curated signatures. 1,000 random 30-gene draws projected through the same top-50 expression-PC basis reproduce the same pan-cancer median *ρ* (−0.73, per-cancer range −0.57 to −0.79; Supplementary Table S16), and in two cancers (BRCA *ρ*_null_ = −0.77, PRAD *ρ*_null_ = −0.79) the random-gene-set null is more negative than the curated empirical value: any 30-gene direction inherits a similar correlation from the leading expression-PC axes. The correlation is nevertheless robust to compositional nuisance: when the null basis is rebuilt as an immune-PC1 ⊕ stromal-PC1 ⊕ proliferation-PC1 anchor block plus the top-47 PCs of the residual expression matrix, the per-cancer correlation becomes *more* negative in 7 of 8 cancers (range −0.76 to −0.91, median −0.86; Supplementary Table S17), ruling out tumor purity, immune infiltrate, and stromal fraction as drivers. Absorption at ExprPC50 is therefore a geometric property of how any 30-gene direction sits in expression-PC space—signatures that recapitulate a few dominant expression-PC axes are strongly absorbed, while signatures whose variance is distributed across many axes retain more residual variance—and this description holds equally for random 30-gene sets. The framework’s curated-vs-random discrimination therefore lives in the absolute-gap finding of the previous paragraph, not in the internal *r*_⊥_-vs-*c* correlation. The curated benchmark’s tier-level distributional structure is consistent with the null-basis geometry described above (Figure 3a). Residual ratios stratify the 17 entries into two distinguishable distributional bands at ExprPC50: the *lower band* contains Tier 2 (immune checkpoint, BRCA DNA repair, cell cycle/CDK; median *r*_⊥_ = 0.059) and Tier 3 (EMT, angiogenesis, apoptosis/BCL2; median 0.072) signatures, whose variance is dominantly absorbed by the top expression-PC basis. The *upper band* contains Tier 1 (TP53, RAS/MAPK, PI3K/AKT/mTOR; median 0.188), Tier 4 (Warburg effect, stemness, NF-*κ*B; median 0.157), and Tier 5 (Fenton metabolic, ceRNA hubs, autophagy, ferroptosis; median 0.202). The across-tier distributional difference is highly significant (Kruskal–Wallis *H* = 63.7, df = 4, *ε*^2^ = 0.50, *p* = 4.9 × 10−13; *n* = 24, 24, 24, 24, 32 for Tiers 1–5), and a complementary two-sided Mann–Whitney test between Tier 1 and all other tiers combined yields *U* = 1830, rank-biserial *r* = 0.47, *p* = 3.9 × 10−4. Crucially, the pairwise Dunn’s post-hoc tests with Benjamini–Hochberg correction (Supplementary Table S4) show that the omnibus signal is driven by Tiers 2 and 3 being significantly lower than Tiers 1, 4, and 5 (all *p*_BH_ ≤ 5 × 10−4), while Tiers 1, 4, and 5 are not pairwise separable (*p*_BH_ = 0.218 for T1 vs T4; *p*_BH_ = 0.978 for T1 vs T5). This tier-level non-separation is consistent with the null-basis geometric description: what distinguishes upper-band from lower-band signatures is how their absorbed variance is spectrally distributed, not which curation tier they belong to, and the same distributional structure is shared by random 30-gene sets projected through the same basis (Supplementary Table S16).

Table 1 makes this concrete. Across the 8 cancers, the top ExprPC50 cell in each cancer spans Tier 1 and Tier 5: TP53 is highest in LUAD (0.364), SKCM (0.442), and HNSC (0.368); autophagy is highest in BRCA (0.408), OV (0.359), and UCEC (0.310); ferroptosis is highest in COAD (0.292) and PRAD (0.223). Warburg effect (Tier 4) is in the upper-band in every cancer (0.211–0.282), and frequently matches or exceeds Tier 1 RAS/MAPK and PI3K/AKT/mTOR in the same cancer. At the same time, the housekeeping control occupies a reproducible middle position (cross-cancer median *r*_⊥_ = 0.149) above the random baseline (0.116) but below the top pathway-specific signatures. That the housekeeping set sits *above* the random baseline is expected rather than paradoxical: a housekeeping gene set is, by construction, biologically coherent and spatially non-random within the transcriptome, so its direction **h** is not a uniform draw from the gene universe and need not match the average absorption of 500 random 30-gene sets. The 50-gene housekeeping control used here is in addition dominated by ribosomal protein genes (40 of 50 entries are RPL∗/RPS∗), and ribosomal protein genes are among the most strongly co-regulated transcripts in bulk cancer expression data; this co-regulation alone is sufficient to concentrate the housekeeping direction onto a small number of dominant expression-PC axes, so the reproducible middle position of this set reflects coherent ribosomal coexpression rather than a true-negative property of “housekeeping biology” as a whole, and the control should be read as a soft non-pathway anchor rather than as an absorption floor. What matters for the framework’s interpretive claims is that the housekeeping set sits *below* every top pathway-specific signature, which it does in every cancer. The co-occurrence of well-validated drivers (TP53) and computationally assembled pathway gene lists (autophagy, ferroptosis) in the same upper band is therefore not a negative result for the framework; it is the framework reporting a structural fact about how these signatures relate to background expression covariance.

The banded architecture generalizes across cancer types: rank correlations of curated-signature residual profiles between cancers range from *ρ* = 0.71 to 0.93 (median 0.82), preserving biologically meaningful context dependence while recapitulating the same lower/upper band decomposition. At the upper-bound ExprPC200 level, Tier 2 falls below Tier 3 largely because the immune checkpoint signature—a Tier 2 member—drops to a near-zero residual ratio by construction when immune PCs are explicitly included in the null-model hierarchy. With the band structure established at the primary operating point, the next question is how often this same pattern recurs across cancers.

### 2.5 Pan-cancer structure at the primary ExprPC50 operating point

At the primary ExprPC50 layer, sub-random behavior is recurrent across every cancer type, although its extent is clearly context dependent rather than universal (Figure 4a). Using the matched random mean as a descriptive anchor, 4 to 10 of the 17 curated benchmark entries fall below baseline in each cancer, with the strongest depletion in OV (10/17) and the weakest in SKCM and HNSC (4/17 each); BRCA and LUAD each show 8/17 entries below baseline, whereas COAD, PRAD, and UCEC fall in between (6/17, 7/17, and 7/17, respectively). Figure 4a provides the complementary inferential summary: after permutation testing against size-matched random controls and Benjamini–Hochberg correction at the pan-cancer curated family level (17 tests per cancer per null-model level; see Methods §4.9), 4–11 curated entries per cancer are classified as significantly below random at *p*_BH_ *<* 0.05. BH correction is applied within each (cancer, null-model) slice (family size = 17), and no cross-cancer or cross-slice correction is applied across the 8 × 8 = 64 (cancer, null-model) slices; this preserves interpretation within each slice as the relevant family for ranking signatures, but family-wise error across the full 64-slice sweep is not controlled and the per-cancer counts above should be read as descriptive prevalence summaries rather than as a unified pan-cancer inferential statement.

**Figure 4:**
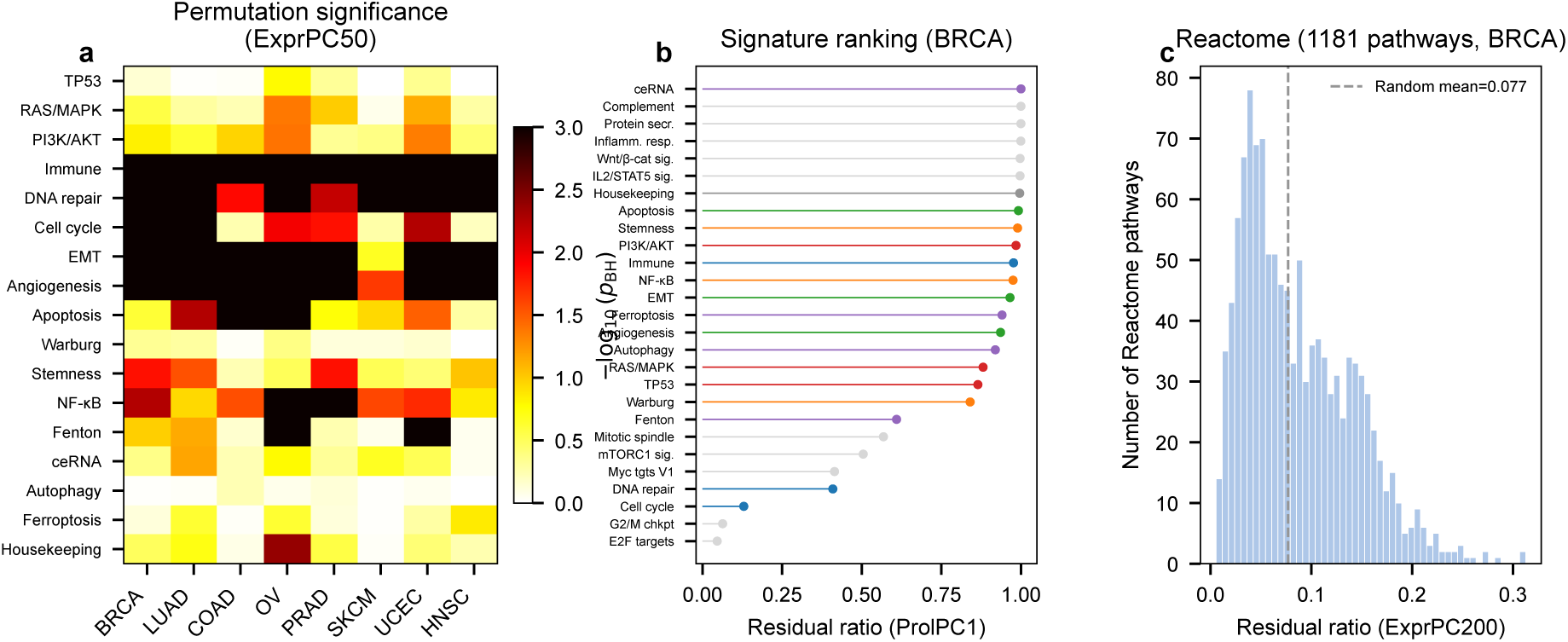
Pan-cancer and public-pathway audit across primary and upper-bound views. (**a**) BH-adjusted permutation significance for falling below size-matched random baselines at ExprPC50 across 8 TCGA cancer types for the 16 curated pathway signatures and the housekeeping control. (**b**) BRCA signature ranking at ProlPC1, combining the curated panel with the five lowest and five highest Hallmark programs. (**c**) Distribution of residual ratios for 1,181 Reactome pathways in BRCA at ExprPC200; the dashed line marks the random mean at this upper-bound sensitivity level.

These descriptive and inferential summaries together indicate that absorption recurs across tumor types but that its extent remains disease dependent. OV, BRCA, and UCEC show broad primary-layer compression of the curated panel, whereas SKCM and HNSC retain a visibly larger fraction of benchmark entries above their random anchor. *Below-random should be read descriptively, not as a quality judgement.* A residual ratio below the matched random baseline indicates that a signature’s variance is more compressed by the null-model basis than an average random gene set of equivalent size; for biologically coherent signatures that track one or two dominant transcriptomic programs (e.g., immune infiltration or cell cycle), this compression is expected rather than pathological. The 4–11 per cancer count therefore quantifies the prevalence of heavy alignment with dominant expression-PC axes in the curated benchmark, not a failure rate, and should be interpreted jointly with the absorption-concentration and overlap analyses presented in §Interpretive boundaries below. The next step is to ask whether related structure persists when the analysis is expanded from the curated panel to larger public pathway collections.

### 2.6 Public pathway collections under upper-bound sensitivity analysis

The expansion to public pathway collections is most informative when read as a sensitivity bound rather than as the manuscript’s default interpretive layer, because it uses a richer null model than the primary operating point (Figure 4b,c). BRCA rankings at ProlPC1 already place curated and Hallmark programs along a shared absorption axis, suggesting that related structure extends beyond the hand-curated benchmark. As an upper-bound sensitivity analysis, ExprPC200 placed 45 of 50 Hallmark gene sets below their matched random baselines in BRCA and 44 of 50 in LUAD. For Reactome, the corresponding counts were 992 of 1,181 pathways in BRCA and 1,000 of 1,181 in LUAD. The BRCA Reactome distribution is shown in Figure 4c, and a full curated-signature ExprPC200 summary is provided in Supplementary Table S3.

Within this upper-bound view, the scale is large but the structure remains heterogeneous. The most strongly absorbed Hallmark categories are immune (median ratio = 0.007, *n* = 7 gene sets), proliferation (0.014, *n* = 5), and DNA/cell-cycle programs (0.013, *n* = 6), whereas tissue-associated programs retain higher residual ratios (median = 0.058, *n* = 6). Category-level analyses likewise show especially strong absorption for ECM/adhesion (*n* = 3, 92% below random), immune (*n* = 7, 87%), and cell-cycle/DNA (*n* = 6, 78%) pathways, whereas subsets of signaling (*n* = 11) and metabolism (*n* = 8) retain more orthogonal variance. Gene-set size is negatively correlated with residual ratio (*ρ* = −0.59, *p <* 0.001), consistent with broader sets sampling more of the dominant coexpression structure, yet the top-ranked sets still include curated driver pathways such as TP53 and PI3K/AKT together with a subset of tissue-associated Hallmark programs. These patterns suggest that rich null models reveal widespread absorption pressure once pathway collections become large and generic enough, while still leaving room for more orthogonal pathway-specific structure in selected programs.

### 2.7 Interpretive boundaries: overlap, absorption mode, and causal ambiguity

Three complementary analyses help bound interpretation of low residual ratios by addressing three common over-readings: that they imply pure artifact, that all absorption is mechanistically equivalent, or that absorption uniquely signals upstream confounding (Figures 5 and 6). First, shared gene membership with canonical pathway databases can import structure into a signature, which means a low residual ratio is not automatically spurious. Among 504 unique genes across the 16 pathway signatures and the housekeeping control, 54 (10.7%) appear in two or more signatures, with the largest overlaps concentrated in metabolism-related signatures and ceRNA network hubs.

**Figure 5:**
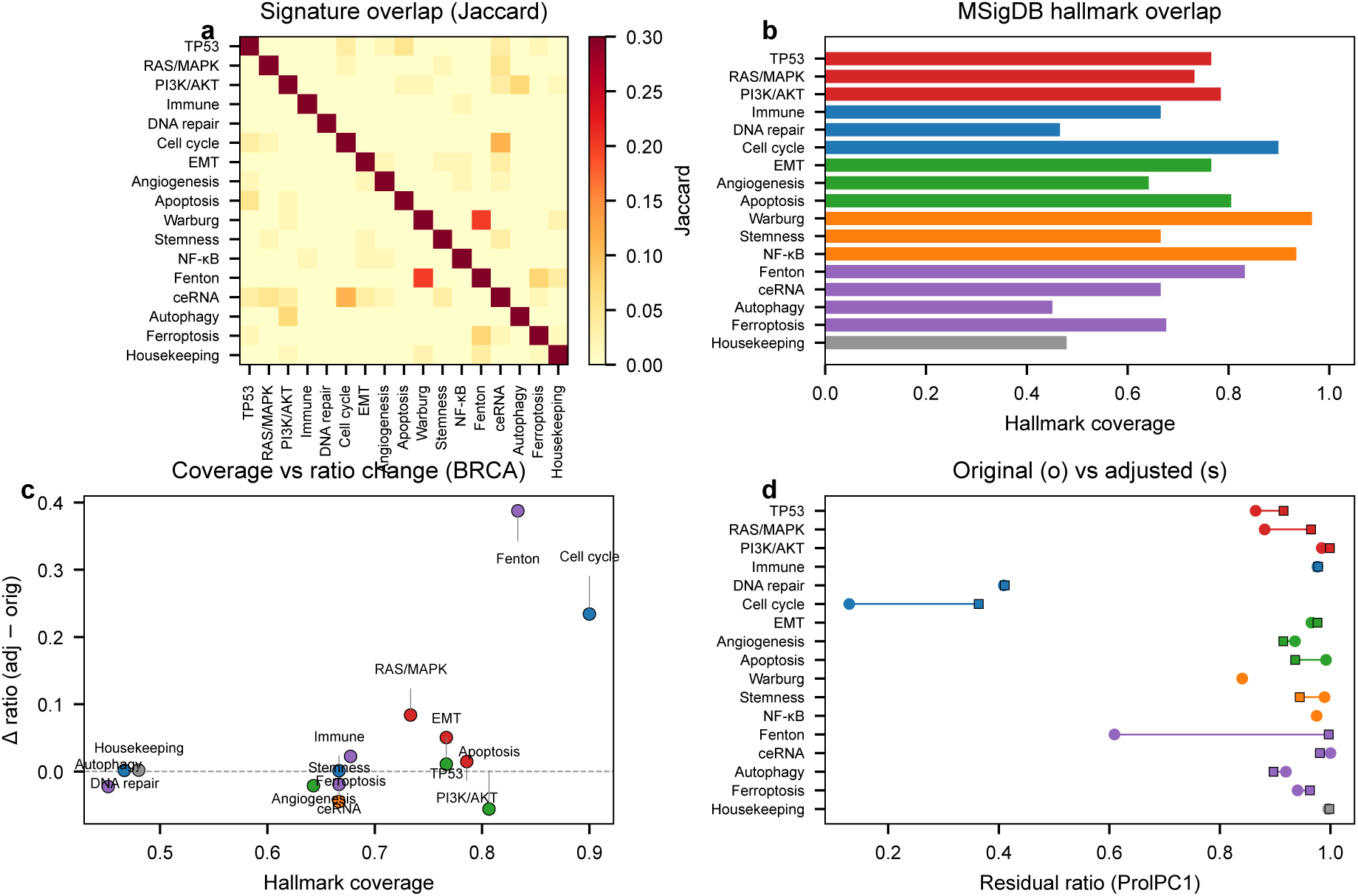
Gene overlap as an interpretive boundary. (**a**) Pairwise Jaccard indices (lower triangle) for the 16 curated pathway signatures and the housekeeping control. (**b**) Fraction of each signature’s genes present in any MSigDB Hallmark set (Hallmark coverage). (**c**) Hallmark gene coverage versus Δ*r*_⊥_ (change after overlap removal) in BRCA (*ρ* = 0.55, *p* = 0.03). (**d**) Original (circles) versus Hallmark-adjusted (squares) residual ratios at ProlPC1; removing shared genes eliminates structured absorption for the most overlap-heavy signatures.

**Figure 6:**
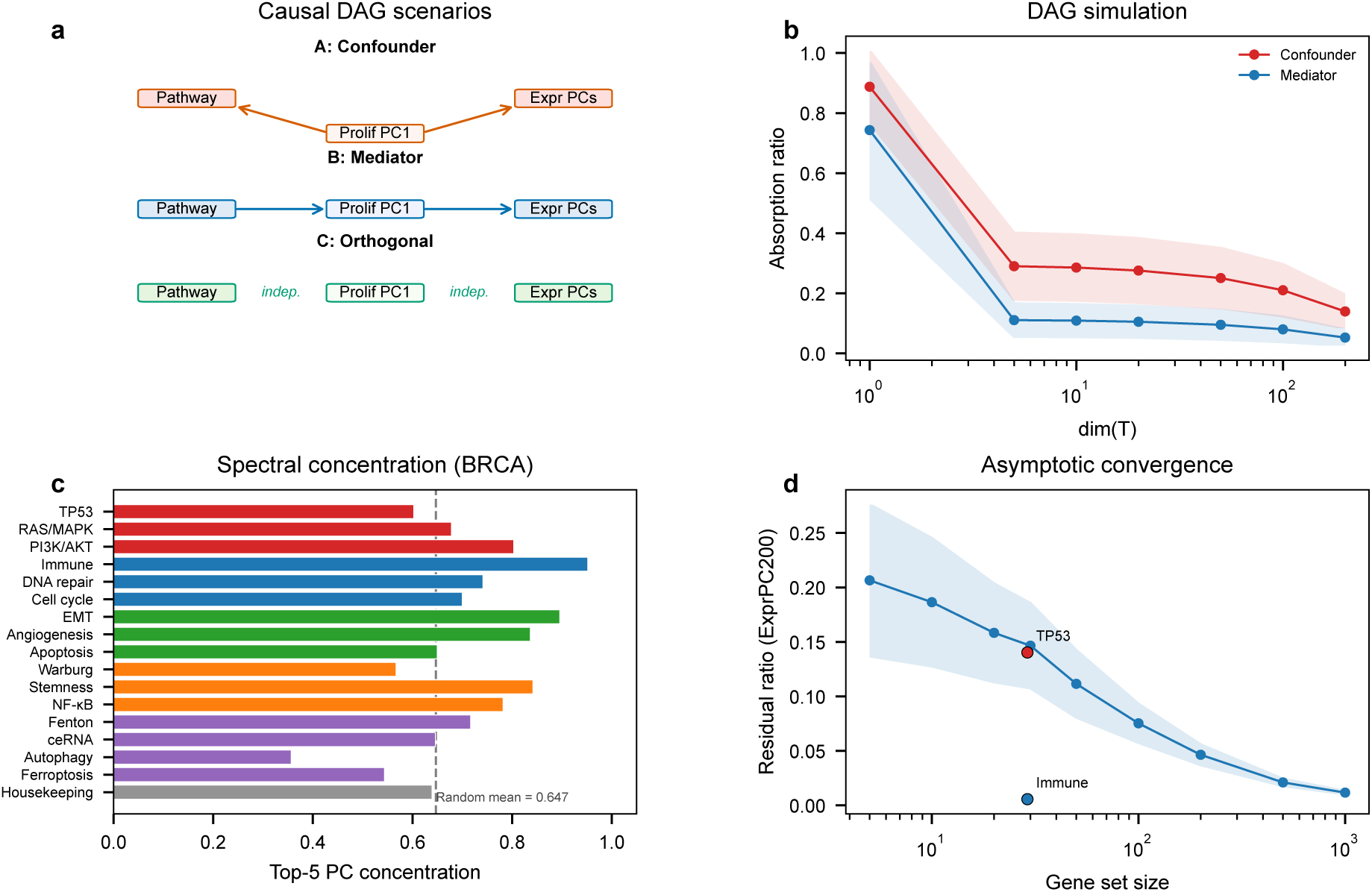
Causal ambiguity and absorption mode as interpretive boundaies. (**a**) Directed acyclic graph (DAG) scenarios used for simulation: *A: Confounder* (latent variable drives both the pathway and the leading expression PCs), *B: Mediator* (pathway drives downstream expression captured by leading PCs), and an orthogonal control (pathway independent of both; not shown graphically but used as the baseline for all gap measurements). (**b**) Absorption trajectories across null-model dimensionality for the confounder-collinear and mediator-driven scenarios in the DAG simulation (mean SD over *n*_rep_ = 100 Monte-Carlo replicates; see Methods §Causal DAG simulation). (**c**) Top-5 absorption concentration 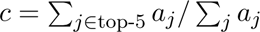 for the curated benchmark in BRCA, showing how much of each signature’s absorbed variance is carried by its five leading absorbing eigencomponents; the dashed line marks the mean *c* = 0.647 of 500 random 30-gene sets (see Methods § Spectral decomposition). Values below the dashed line indicate more diffuse absorption than random. (**d**) Asymptotic convergence analysis relating gene-set size to residual ratio at ExprPC200, with TP53 and immune checkpoint highlighted.

Hallmark coverage ranges from 45% (Autophagy) to 83% (metabolism-related signatures), and in BRCA higher Hallmark coverage is associated with larger overlap-adjusted shifts in residual ratio (Spearman *ρ* = 0.55, *p* = 0.03). In the strongest example, removing Hallmark-overlapping genes raises a metabolism signature in BRCA from *r*_⊥_ = 0.61 to near 1.0, whereas TP53 remains high after removal (0.87 → 0.92). The same trend is absent in LUAD (*ρ* = 0.06), consistent with the view that no single overlap mechanism explains all low-residual signatures.

Second, not all absorption looks the same once its spectral distribution is examined. The primary concentration statistic we report is the *inverse participation ratio* (IPR, Methods § 4.7), 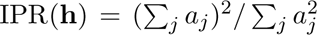, which measures the effective number of eigencomponents that participate in absorbing the signature direction **h**; for readability we also quote the top-5 concentration 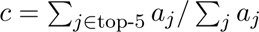. Computed across all 17 curated benchmark entries in each of the 8 TCGA cancers (Supplementary Table S10), concentration tracks absorption at panel scale in every cancer: the Spearman rank correlation between *c* and the ExprPC50 residual ratio is negative in every cancer (range −0.59 in PRAD to −0.89 in SKCM, median −0.71; all 8 significant at *p <* 0.05), and the correlation between IPR and ExprPC50 residual ratio is positive in every cancer (range +0.51 in BRCA to +0.86 in SKCM, median +0.62). As established in §Curated-panel synthesis, these panel-scale correlations are descriptive geometric properties of the null-model coordinate system rather than specific properties of curated biology—1,000 random 30-gene draws per cancer reproduce the same pan-cancer median *ρ* (Supplementary Table S16)—and we report them here as a consistency check on how any gene-set direction sits in expression-PC space: signatures whose absorbed variance is concentrated on a small number of leading expression-PC axes are the most strongly absorbed (lowest *r*_⊥_), whereas signatures that distribute absorption across many axes retain the largest residual variance. The two exemplars illustrate the tails of this distribution. Immune checkpoint in BRCA concentrates *c* = 0.96 of its absorbed variance into its top-5 PCs with an effective rank of only IPR ≈ 4.0, reflecting a strongly few-axes pattern. TP53 absorption is more diffuse (*c* = 0.75, IPR ≈ 6.5 in BRCA) and autophagy still more diffuse (*c* = 0.50, IPR ≈ 14.1 in BRCA, above the random-baseline mean IPR = 5.1 ± 2.9). Low residual ratio therefore does not describe a single mechanism: at panel scale the dominant mode is concentrated alignment with a few dominant axes (as in immune checkpoint, EMT, angiogenesis, cell cycle/CDK, stemness), while a minority of signatures are both highly absorbed and spectrally diffuse (autophagy, ferroptosis, housekeeping), indicating that absorption at ExprPC50 is *primarily but not exclusively* explained by few-axis alignment with dominant transcriptomic programs. This is a structural result across the full panel of 17 signatures in all 8 cancers, not an interpolation from two exemplar signatures.

Third, projection-based null models absorb both upstream confounding and downstream pathway consequences, so absorption cannot be equated with confounding alone. To quantify this ambiguity, we conducted causal simulations under a known directed acyclic graph (DAG) with three scenarios: (i) a gene set whose variance is driven by an upstream confounder (confounder-collinear), (ii) a gene set that causally drives downstream transcriptomic effects captured by expression PCs (mediator-driven), and (iii) a gene set orthogonal to both (Figure 6a). At low null-model dimensionality (ExprPC5), with *n*_rep_ = 100 Monte-Carlo replicates per scenario, the confounder-collinear and mediator-driven scenarios are distinguishable: the former retains *r*_⊥_ = 0.273 ± 0.136 (95% CI on the mean [0.246, 0.300]) and the latter *r*_⊥_ = 0.128 ± 0.100 (CI [0.108, 0.148]), yielding a mean gap of 0.144 (95% CI [0.111, 0.178]; Figure 6b). This gap narrows at higher dimensionality, contracting to ≈ 0.07 at ExprPC200.

A 27-configuration sensitivity sweep around the base setting (*N* = 500, *G* = 5,000, *k*_sig_ = 30) independently varies confounder strength, pathway propagation strength, residual noise scale, pathway gene-set size, and number of downstream genes (Supplementary Table S6). 24 of 27 configurations (89%, Clopper–Pearson 95% CI 0.71–0.98) preserve distinguishability at ExprPC5. Within the simulated range *k* ∈ {5, 10, 20, 50, 100, 200}, the residual ratio of the confounder-collinear scenario contracts monotonically from 0.29 at *k* = 5 to 0.14 at *k* = 200 (equivalently, the absorbed fraction grows from 0.71 to 0.86; Figure 6d). Because direct Monte-Carlo simulation is not feasible once *k > N* − 2 under the base configuration, we make no claim beyond *k* = 200 and treat ExprPC200 as the empirical upper bound on absorption for this simulation.

#### Numerical overlap between simulated and real signatures

At ExprPC50 the confounder-collinear scenario retains *r*_⊥_ = 0.233 (95% CI [0.210, 0.256]; comparable to real Tier 1 validated drivers: cross-cancer median 0.188; TP53 cross-cancer median 0.283), whereas the mediator-driven scenario retains *r*_⊥_ = 0.110 (CI [0.093, 0.127]; comparable to Tier 2–3 real signatures in the lower band of the curated panel). A single residual-ratio reading at any one operating point is therefore *not* a sufficient discriminator between confounder-driven and driver-driven transcriptomic signals: a non-zero residual ratio at ExprPC50 should not by itself be read as evidence that a signature is confounder-independent, and a low residual ratio should not by itself be read as evidence that a signature is confounded. In effect, absorption at ExprPC50 is a *geometric* property of how the signature direction lies with respect to the leading expression-PC axes, and the overlap with Tier 1 values is the mechanical consequence. For translational readers, two consequences follow. First, the confounder–mediator gap narrows by construction as *k* grows: 0.144 at ExprPC5 contracts to ≈ 0.07 by ExprPC200, so confounder/driver discrimination is not accessible at the primary ExprPC50 operating point under this framework’s current linear-projection formalism. Second, the numerical overlap between the confounder-collinear simulation and TP53’s cross-cancer median residual ratio is a statement about the residual-ratio metric at *k* = 50, not about the biological status of TP53-pathway signatures: a reader should not interpret *r*_⊥_(TP53) ≈ 0.28 as evidence that TP53 signatures are confounder-driven, nor as evidence that they are confounder-independent. This is a bound on the framework’s interpretive scope, not on TP53 biology.

#### The trajectory shape is empirically stable

The claim that the trajectory, rather than any single *r*_⊥_(*k*_0_), is the framework’s load-bearing signal rests on an empirical stability check. Across 200 sample-level fixed-basis bootstrap resamples of the BRCA cohort (Supplementary Table S14; Methods §Statistical analysis), each of the 17 curated benchmark entries yields a family of trajectory vectors *r*_⊥_(*k*) at *k* ∈ {1, 5, 10, 20, 50, 100, 200} whose mean pairwise Pearson correlation across resamples is 0.999 (minimum 0.998, maximum 1.000). The *shape* of the trajectory is therefore effectively deterministic at the sample level. Individual cell values at a single null-model dimensionality under *B* = 1,000 fixed-basis resamples are also tight: the BRCA median 95% CI half-width at ExprPC50 is 0.019 and the LUAD median is 0.024 (Supplementary Table S5), with an overall range of ∼ 0.002 (immune checkpoint, whose absolute value is itself ∼ 0.02) to ∼ 0.05 (autophagy and TP53 in LUAD). Both the shape and the cells are therefore bootstrap-reliable, with the shape being effectively invariant and the individual cell values tight enough for moderate contrasts (differences ≳ 0.05) but not for fine distinctions within the upper band. This gap—an essentially deterministic shape and cell-level uncertainty on the order of 0.02–0.05—is the statistical justification for reading the trajectory shape as the framework’s primary signal and treating any single *r*_⊥_(*k*_0_) difference smaller than ∼ 0.05 as uncertain. Combined with the absorption concentration (Figure 6c), gene overlap with canonical programs (Figure 5), and matched-random percentile rank, the trajectory shape is the framework’s primary reporting object: residual-ratio auditing provides a bootstrap-stable description of how a signature’s variance relates to a modeled background subspace, not a causal verdict or a single-number confounder detector.

## 3 Discussion

### 3.1 What the residual ratio measures

The residual ratio quantifies how much of a gene signature’s sample-space variance remains orthogonal to a specified background-expression model. It is therefore a property of three objects jointly: the signature, the data, and the null model. Changing any of these changes the ratio. Under the linear-projection formalism used here (Methods § 4.7), *r*_⊥_ is a purely *geometric* property: given a signature direction **h** in sample space and a null-model basis **Q** comprising the top expression PCs, *r*_⊥_ = ^‖^**h** − **QQ**𝖳**h**‖2 is the squared length of the component of **h** orthogonal to col(**Q**). This is not a biological-validation claim: a signature can have a high residual ratio because it captures genuinely orthogonal biology, because it is noisy, or because it is aligned with a latent factor that the null model does not happen to span. Disambiguating these possibilities requires the trajectory shape, the matched-random baseline gap, and the absorption concentration, read jointly. A low residual ratio does not imply that a signature lacks biological relevance; it indicates that, in the dataset and null model under study, the signature’s variation is not easily distinguished from the modeled background structure.

We define “background expression structure” operationally as the subspace spanned by the null model’s covariates. At ExprPC50, this subspace captures 61.6% of total transcriptomic variance in BRCA (log-transformed, median-filtered expression matrix, *G* = 16,680 genes, *N* = 1,082 samples); at ExprPC200, 79.2%. Per-cancer cumulative variance at both operating points is reported in Supplementary Table S9. For interpretation, the most informative quantity is often the *trajectory* of residual ratios across the hierarchy, together with comparison to matched random baselines, rather than the value at any single null-model level. ExprPC50 provides a practical operating point, whereas ExprPC200 is best treated as an upper bound on absorption.

### 3.2 What the framework does not establish

Residual-ratio auditing addresses a question that is complementary to, rather than competitive with, existing pathway-scoring methods such as GSVA, ssGSEA, singscore, and PROGENy [73–76]. These methods estimate *how active* a pathway appears in each sample; our framework asks *how much of that variation is distinguishable from modeled background transcriptomic covariance*. A pathway may score highly while still having a low residual ratio if its variation largely recapitulates proliferation-associated or immune-associated structure.

The framework also does not provide a causal adjudication, and residual ratio at a single operating point is *not* a reliable confounder detector. Expression PCs can include both confounders and downstream mediators of pathway activity, as illustrated by the causal simulations in Section 2.7: in particular, a simulated signature entirely driven by a latent confounder retains *r*_⊥_ ≈ 0.25 at ExprPC50 (Results § Interpretive boundaries), a value comparable to real Tier 1 validated drivers. For a mutationally driven pathway such as TP53, conditioning on expression PCs may additionally remove some of the pathway’s own downstream consequences alongside nuisance structure. The trajectory of residual ratios across the null-model hierarchy, combined with the matched-random baseline gap and overlap assessment, is therefore the framework’s load-bearing signal, and experimental perturbation remains essential for establishing pathway-specific causation [60].

To position residual-ratio auditing directly against the closest prior-art signature-quality framework, we reproduced a Berglund-style “uniqueness” metric [83], operationalized as 1 − max*_j_ a_j_*, where *a_j_* is the squared projection of the signature direction onto the *j*-th leading expression PC; high uniqueness means that no single global PC explains a large share of the signature. We then computed Spearman rank correlations between Berglund uniqueness and the ExprPC50 residual ratios reported here at three scales: the 17-entry curated benchmark, all 50 MSigDB Hallmark gene sets, and all 1,361 Reactome pathways in the Enrichr Reactome library’s most recent snapshot (filtered to 15–500 genes; the 180-pathway difference from the 1,181 Reactome 2022 set used for our primary audit under the identical 15–500 gene filter reflects content growth in the Reactome 2022→2024 window rather than any methodological change, and we report the Berglund head-to-head against the more recent snapshot because that is the Enrichr default), each in BRCA and LUAD. Per-entry values and full head-to-head tables are provided in Supplementary Tables S7 and S11.

At the 17-entry curated benchmark the two metrics show moderate agreement (BRCA Spearman *ρ* = 0.57, Fisher-*z* 95% CI 0.12–0.82; LUAD *ρ* = 0.62, CI 0.19–0.85). This *n* = 17 agreement is too small to distinguish a “noisy-but-redundant” relationship from a genuinely complementary one—the curated-scale CI alone spans from weak to strong. Extending the comparison to the Hallmark scale (*n* = 50) tightens the CI substantially (BRCA *ρ* = 0.39, CI 0.12–0.60, *p* = 5.2 × 10−3; LUAD *ρ* = 0.61, CI 0.40–0.76, *p* = 2.5 × 10−6), and extending it to the Reactome scale (*n* = 1,361) tightens the CI to an order of magnitude narrower (BRCA *ρ* = 0.55, CI 0.51–0.59, *p <* 10−100; LUAD *ρ* = 0.67, CI 0.64–0.70, *p <* 10−100). At all three scales the correlation is positive, moderate-to-strong, and the two diagnostics disagree most visibly at signatures whose absorption is dominated by *one to two* specific axes: the immune checkpoint signature is ranked last by the residual ratio at ExprPC50 (where the full 50-dimensional null-model subspace is projected out) but only moderately low by Berglund uniqueness (where 1 − max*_j_ a_j_* is insensitive to absorption distributed across two or three near-top PCs). The conceptual difference is clean and now quantified at three scales: Berglund uniqueness asks whether a signature score is redundant with any *single* global PC, whereas residual-ratio auditing asks whether the signature direction is orthogonal to the *subspace* spanned by an explicitly chosen set of null-model covariates, and supports a progressive hierarchy of null-model richness. Because the Hallmark and Reactome Spearman CIs at full scale exclude zero by many orders of magnitude, the two diagnostics are neither perfectly redundant nor independent; they capture closely related but non-identical aspects of the same absorbed-variance decomposition.

### 3.3 How to interpret low residual ratio

A low residual ratio can arise for several distinct reasons. First, a signature may align strongly with dominant transcriptomic axes such as proliferation, immune infiltration, or stromal composition, in which case variance is lost early under biologically interpretable null models. Second, gene overlap with canonical programs can transfer residual structure into signatures that borrow genes from well-characterized pathways, although such overlap can also reflect convergent curation rather than simple contamination. Third, larger gene sets sample more broadly across coexpression modules and therefore more closely approximate the transcriptome-wide mean direction, which is by construction aligned with leading expression PCs.

Absorption concentration provides a useful secondary diagnostic in this setting. The parameter-free inverse participation ratio IPR (Methods § 4.7) reports the *effective number* of expression-PC axes that together absorb the signature: IPR close to 1 indicates single-axis absorption, whereas IPR approaching *p* indicates uniform spreading across the full null-model basis. Across the 17 curated BRCA entries, the immune checkpoint signature shows IPR ≈ 4.0 with top-5 concentration *c* = 0.96 (absorption dominated by a small immune-aligned subspace), TP53 shows IPR ≈ 6.5 with *c* = 0.75 (absorption distributed across more axes), and autophagy shows an even more diffuse IPR ≈ 14.1 with *c* = 0.50, below the random baseline (*c* = 0.65 ± 0.11; random 30-gene IPR ≈ 5.1 ± 2.9). As reported in Results §Curated-panel synthesis, the panel-wide Spearman correlation between *r*_⊥_ at ExprPC50 and *c* is negative in every cancer (median −0.71 across 8 cancers); this is a descriptive geometric property of the null-model coordinate system that random 30-gene sets reproduce at the same pan-cancer median (Supplementary Table S16), and is reported here as a consistency check on how any signature direction sits in expression-PC space rather than as a biological discovery specific to the curated panel.

#### Read the trajectory shape, not a single cell in isolation

When comparing two signatures on a per-cell basis, the trajectory shape {*r*_⊥_(*k*)} provides the most statistically reliable comparison. Across 200 sample-level bootstrap resamples the trajectory-shape vectors of the 17 curated entries in BRCA are invariant at mean pairwise Pearson *ρ* = 0.999 (Supplementary Table S14), and individual cell-level *r*_⊥_ values at ExprPC50 under *B* = 1,000 fixed-basis resamples have 95% bootstrap CI half-widths of approximately 0.019 in BRCA and 0.024 in LUAD (overall range ∼ 0.002–0.053). Two cells in Table 1 that differ by ≲ 0.05 may therefore lie within mutual bootstrap uncertainty, while differences of ≳ 0.08 are generally significant at the cell level in BRCA. Summary statistics over the panel—distributional bands, tier medians, and the curated-vs-random magnitude gap—are consequently tighter than individual cell-level contrasts and are the appropriate currency for pathway-level interpretive claims. Low residual ratio does not by itself imply a lack of practical value: a proliferation-aligned signature may still be clinically useful precisely because it tracks a strong prognostic program. The framework evaluates pathway-level interpretation claims, not clinical utility per se.

### 3.4 How to use the framework in practice

Based on these analyses, we suggest a compact reporting workflow for studies that make pathway-level interpretation claims from transcriptomic gene signatures (Table 2). We do not view this workflow as a binary pass/fail standard. Rather, it is intended as a practical starting point to help authors report whether a signature remains orthogonal to interpretable background structure, how it compares with matched random baselines, whether overlap with canonical databases materially changes the result, and whether conclusions remain stable across alternative scoring schemes or datasets.

**Table 2:** Suggested reporting elements for transcriptomic gene-signature audits. A practical workflow for studies making pathway-level interpretation claims from transcriptomic signatures.

| # | Audit item | Description |
| --- | --- | --- |
| 1 | Trajectory $r_{\perp}(k)$ | Report the full trajectory of residual ratios across the null-model hierarchy (e.g., ProlPC1, ExprPC5, ExprPC20, ExprPC50, ExprPC100, ExprPC200) rather than the value at any single $k$. The trajectory *shape* is the bootstrap-stable quantity; single operating-point values should be reported alongside sample-level bootstrap 95% CIs. |
| 2 | Spectral concentration | Report the top-5 absorption concentration $c$ and/or the parameter-free inverse participation ratio IPR alongside $r_{\perp}$. These diagnostics describe how each signature’s absorbed variance is spectrally distributed (few-axis concentrated vs. diffuse across many PCs), independent of its absolute $r_{\perp}$ value. |
| 3 | Random baseline | Compare against size-matched random gene sets ($n \geq 500$) from the same expression matrix; report percentile rank in terms of both $r_{\perp}$ and $c$. |
| 4 | Overlap profile | Report Jaccard overlap with MSigDB Hallmark, Reactome, and other canonical pathway databases. |
| 5 | Overlap-adjusted ratio | Remove genes shared with canonical databases and recompute $r_{\perp}$; report the change. |
| 6 | Scoring robustness | Verify that conclusions hold under $\geq 2$ alternative scoring methods (e.g., z-score, rank-based). |
| 7 | Cross-dataset consistency | Report the trajectory $r_{\perp}(k)$ across $\geq 2$ independent cohorts or cancer types. |

We deliberately omit fixed thresholds because the appropriate benchmark depends on the null model, cancer type, gene-set size, and research question. In practice, we recommend reading the trajectory shape together with the gap between each signature’s *r*_⊥_ and its matched random baseline as the two statistically reliable outputs of the framework, while treating any single operating-point value as a coarser summary whose bootstrap uncertainty should be explicitly quoted. When residual ratios fall to only 4–5% under very rich null models, the remaining orthogonal signal becomes correspondingly small and the trajectory itself carries most of the interpretive information.

### 3.5 Limitations and scope

Our framework has several limitations. First, at the ExprPC200 level, the null model’s principal components are computed from all genes, including those in the signature being tested. We verified that excluding signature genes from the PC computation changes residual ratios by less than 0.02 for key signatures in BRCA (TP53: +0.016; Fenton: +0.003; BRCA repair: +0.002). Second, the residual decomposition is linear; nonlinear confounding relationships (e.g., threshold effects, epistasis) are not captured. Third, the choice of null-model dimensionality is consequential: we use ExprPC50 as a consensus operating point supported by cross-validated selection (Methods §Cross-validated dimensionality selection), but alternative criteria could yield different recommendations. Fourth, we use uniform gene weights; weighted signatures may show different residual profiles. Fifth, our analysis is based entirely on bulk RNA-seq (TCGA PanCancer Atlas, 8 primary cancer types) and microarray (METABRIC) data; bulk measurements confound cell-type-specific signals with compositional effects, and the framework’s empirical claims should not be transferred to single-cell RNA-seq, single-nucleus RNA-seq, or spatial transcriptomics without separate evaluation—linear PCA-based null models rest on assumptions (approximately Gaussian sample-space covariance, limited dropout, no per-cell count-depth heterogeneity) that do not hold in the same form for sparse single-cell or spatially correlated data, and residualization strategies appropriate to those modalities (GLM-PCA, randomized SVD on sparse matrices, block bootstrap for Visium/Xenium) are out of scope for this work. Sixth, the causal DAG simulation uses a simplified generative model, whereas real transcriptomic causal structures are far more complex.

For these reasons, we view residual-ratio auditing as a complementary interpretive layer rather than a standalone decision rule. A high residual ratio indicates that a signature’s variance is distinguishable from the modeled background structure in the dataset under study—not proof of causation; a low residual ratio indicates strong alignment with modeled covariance—not automatic lack of usefulness. The framework is most informative when used together with explicit null-model choice, matched baselines, overlap assessment, and independent biological knowledge.

## 4 Methods

We designed the benchmark to quantify how much of each transcriptomic gene signature’s variance remains orthogonal to progressively richer null models of background expression structure, across curated and public pathway collections, while preserving interpretable links between the null model and the resulting residual ratio. The openly available implementation, benchmark outputs, and gene signature definitions are described below and in the accompanying repository.

### 4.1 Data sources

We obtained RNA-Seq V2 RSEM gene expression data for ten TCGA cancer types from the PanCancer Atlas [1] via the cBioPortal data repository [4, 5], which provide open-access, version-stable mRNA expression matrices for each cancer study under standard PanCancer Atlas identifiers; these resources complement related genomic collections such as the International Cancer Genome Consortium [71] and the Human Protein Atlas [72]. Cancer types with fewer than 200 samples (GBM and PAAD) were excluded from the primary analysis because the ExprPC200 upper-bound null model requires *N* ≥ 201+2 samples to support a rank-201 orthogonal projection without strong truncation of the principal-component basis (§Null model hierarchy describes the truncation rule applied to cancer types just above the cutoff); a supplementary ExprPC50 analysis that includes both cohorts is reported in Supplementary Table S8 and shows the same qualitative tier gradient, confirming that the exclusion does not alter the primary interpretation. Eight cancer types were retained for the primary analysis (Table 1): BRCA (*n* = 1,082), LUAD (*n* = 510), COAD (*n* = 592), OV (*n* = 300), PRAD (*n* = 493), SKCM (*n* = 443), UCEC (*n* = 527), and HNSC (*n* = 515), totalling 4,462 samples. Expression values were log_2_(RSEM+1)-transformed, and genes with median expression below 1.0 across samples were removed, yielding approximately 16,500–16,900 genes per cancer type. No additional batch correction beyond the PanCancer Atlas release-level normalization was applied; the framework’s numerical outputs depend on the batch-correction state of the input expression matrix, and we recommend that downstream users report whether and how their input was batch-corrected (see Discussion §3.5). Clinical data for BRCA (age, AJCC stage, fraction genome altered) were obtained from the same source. For external validation, we obtained microarray expression data from the METABRIC breast cancer cohort (*n* = 1,980) from cBioPortal. The exact study identifiers, file names, and download dates for every cancer cohort used in the primary and supplementary analyses are listed in the Declarations section (Availability of data and materials).

### 4.2 Residual decomposition

For a gene expression matrix **Y** ∈ R*^N^*^×*G*^ (*N* samples, *G* genes) and a gene signature with weight vector **w** ∈ R*^G^*, we define the signature’s sample-space direction:

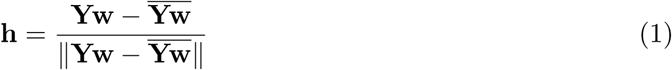

where the overline denotes the sample mean and the denominator normalizes to unit norm. Given a null model design matrix **X** ∈ R*^N^*^×^*^p^* (including intercept), we compute its orthonormal column basis **Q** ∈ R*^N^*^×^*^p^* via QR decomposition. The orthogonal projection onto the null model subspace *T* = col(**X**) is:

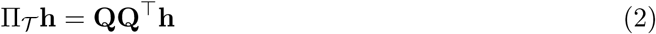

The residual is **R** = **h** − Π_𝓨_ **h**, and the *residual ratio* is:

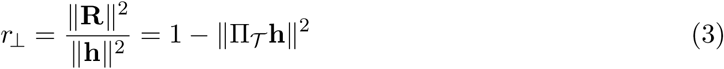

which lies in [0, 1]. A ratio of 0 indicates that **h** ∈ *T* (the signature is a linear combination of confounders); a ratio of 1 indicates complete orthogonality. This quantity is equivalent to 1 − *R*^2^ from multiple linear regression of the signature score on the null model covariates.

### 4.3 Null model hierarchy

We constructed eight progressively enriched null models:

1. **Intercept only** (dim = 1): intercept-only baseline.
2. **Prolif PC1** (dim = 2): first principal component of 50 proliferation marker genes (Table S2) [85].
3. **Prolif PC1–10** (dim = 11): top 10 PCs of proliferation genes.
4. **Prolif + Clinical** (dim ≈ 24): top 20 proliferation PCs plus age, stage, and fraction genome altered (when available).
5. **Full biological** (dim ≈ 34): above plus top 10 PCs of immune marker genes.
6. **ExprPC50** (dim = 51): top 50 PCs of the full expression matrix (primary interpretive operating point; strict CV optimum in BRCA, CV-admissible in 6 of 7 other primary cancers; see §Cross-validated dimensionality selection).
7. **ExprPC100** (dim = 101): top 100 expression PCs.
8. **ExprPC200** (dim = 201): top 200 expression PCs (capturing ∼79% of total variance in BRCA; upper bound).

For cancer types with sample size *N <* 200 + 2, the ExprPC200 level was capped at *N* − 2 components.

### 4.4 Gene signature curation

We benchmarked 16 curated pathway signatures organized into five validation tiers, together with a housekeeping control and random-gene-set baselines. We emphasize that the tier labels are *descriptive, domain-knowledge annotations* assigned by the authors before analysis to organize how each gene list was assembled and validated in prior literature, not pre-registered hypothesis classes; the framework’s primary empirical claim (the magnitude-gap finding of §Curated-panel synthesis) does not depend on the tier assignment, and tier-level distributional statistics in the main text are presented as descriptive summaries of the panel rather than as hypothesis tests against the tier labels.

- **Tier 1 — Well-validated drivers** (3 signatures): TP53 pathway [27], RAS/MAPK [28], PI3K/AKT/mTOR [29]. All have FDA-approved targeted therapies and decades of experimental validation.
- **Tier 2 — Clinically validated** (3 signatures): Immune checkpoint [30], BRCA DNA repair [31], Cell cycle/CDK [32]. All have approved clinical interventions.
- **Tier 3 — Experimentally validated** (3 signatures): EMT [33], Angiogenesis [34], Apoptosis/BCL2 [35]. Functional validation exists but clinical translation is limited.
- **Tier 4 — Partial validation** (3 signatures): Warburg effect [36], Stemness [37], NF-*κ*B inflammatory [38]. Partial experimental support.
- **Tier 5 — Computationally assembled gene lists** (4 signatures): Fenton metabolic [43, 44], ceRNA network hubs, Autophagy (TCGA-derived), Ferroptosis. We emphasize that this tier classification refers to the *specific gene lists used here*, which were assembled computationally, not to the underlying biological processes. Both ferroptosis [39] and autophagy [42] have extensive experimental validation as biological phenomena; our Tier 5 designation indicates only that these particular transcriptomic gene lists have not been independently validated as faithful proxies for pathway activity in bulk tumor data.
- **Controls**: Housekeeping genes (50 ribosomal proteins and metabolic enzymes) and random 100-gene sets (*n* = 200 per null model level).

Each pathway signature contains 28–66 genes (median 30), and the housekeeping control contains 50 genes. Gene weights were uniform: 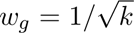 for each of *k* genes present in the expression matrix, and zero otherwise. Gene lists are provided in Table S1.

### 4.5 MSigDB Hallmark and Reactome pathway audits

We audited all 50 MSigDB Hallmark 2020 gene sets [52] and 1,181 Reactome 2022 pathways (filtered to 15–500 genes) identically to the curated signatures. Per-gene-set size-matched random baselines (200 replicates for Hallmark; 100 for Reactome) were used for percentile rank computation.

### 4.6 Causal DAG simulation

To distinguish confounder-collinear from mediator-driven absorption, we simulated gene expression data under a known DAG with three gene-set scenarios. The generative model specifies: (i) a latent confounder *Z* ∼ N(0*, I_k_Z*) with *k_Z_* = 5 that drives both the confounder-collinear gene set and background expression; (ii) a causal pathway whose gene-set score drives downstream expression changes captured by *n*_downstream_ downstream genes and therefore by the leading expression PCs; and (iii) an orthogonal gene set independent of both. The base configuration used *N* = 500 samples, *G* = 5,000 genes, pathway gene-set size *k*_sig_ = 30, *n*_downstream_ = 500, confounder strength 0.8, propagation strength 1.5, and residual noise scale 1.0. For each scenario, we computed the residual ratio across null-model dimensionalities *k* ∈ {1, 5, 10, 20, 50, 100, 200} from *n*_rep_ = 100 independent Monte-Carlo replicates and measured the gap between confounder-collinear and mediator-driven residual ratios as a function of *k*. We report gap 95% confidence intervals using a Gaussian approximation with standard errors 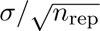 pooled over the two scenarios.

#### Sensitivity sweep

We ran a 27-configuration one-at-a-time sensitivity sweep around the base configuration by independently varying each of five parameters (confounder strength over 6 values; propagation strength over 6 values; noise scale over 5 values; pathway gene-set size *k*_sig_ over 5 values; number of downstream genes *n*_downstream_ over 5 values), totalling 27 configurations including the base. For each configuration we re-ran the DAG scenarios and recorded whether the confounder-collinear vs mediator-driven gap at *k* = 5 remained positive and non-overlapping. 24 of 27 configurations (89%, Clopper–Pearson 95% CI 0.71–0.98) preserved distinguishability at ExprPC5; the full sweep is provided in Supplementary Table S6.

#### Finite-*k* regime only

Because direct Monte-Carlo simulation is not feasible once *k > N* − 2 under our base configuration (*N* = 500), all reported DAG simulation results are restricted to *k* ∈ {1, 5, 10, 20, 50, 100, 200}; we make no asymptotic claim beyond *k* = 200 and treat ExprPC200 as the empirical upper bound on absorption for the simulation. Values at larger *k* would require either scaling *N* or deriving a rank-deficient closed-form, neither of which is required for the interpretive claims reported in the main text.

### 4.7 Spectral decomposition of absorbed variance

For a signature direction **h** (unit-norm by construction) and a null-model basis **Q** = [**q**_1_, …, **q***_p_*] whose columns are the orthonormal ExprPC directions computed from **Y**, we decompose the absorbed variance into per-eigencomponent contributions

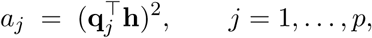

so that the total absorbed fraction is 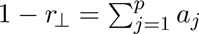 and ‖**h**‖2 = 1. Because each *a_j_* ≥ 0, the normalized absorbed spectrum 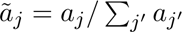 is a probability distribution over eigencomponents, which makes the *shape* of the spectrum—not just its total mass—meaningful.

#### Primary concentration statistic: inverse participation ratio (IPR)

As the primary, parameter-free measure of how broadly the absorbed variance is distributed across eigencomponents, we report the inverse participation ratio

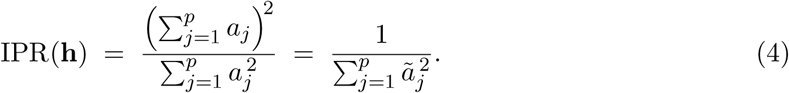

IPR has units of eigencomponents and admits a clean interpretation: IPR = 1 when one eigencomponent carries all of the absorbed variance, IPR = *p* when absorption is uniformly distributed across all *p* eigencomponents of the null-model basis, and intermediate values give the *effective number* of eigencomponents that participate in absorbing **h**. A low-IPR signature (few-axis absorption) is aligned with a small number of dominant expression-PC axes; a high-IPR signature spreads its absorption across many axes, even if its total absorbed mass is similar. Because IPR depends only on the normalized absorption spectrum *ã_j_*, it is invariant to rescaling of **h** and does not require an operating-point-dependent cutoff.

#### Auxiliary concentration statistics

For readability and cross-comparison with prior work, we also report the *top-5 absorption concentration* 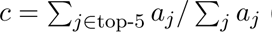 (the fraction of absorbed variance carried by the five eigencomponents with the largest *a_j_* values) and the *single-leading-axis contribution* 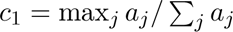. Both are auxiliary summaries of the same normalized spectrum *ã_j_*: *c* is a fixed-*k* truncation and *c*_1_ is the *ℓ*_∞_ norm of *ã_j_*. Neither is parameter-free, and both are degenerate when *p <* 5; we therefore treat IPR as the primary concentration statistic and *c, c*_1_ as readability aids. As an information-theoretic cross-check we additionally report the Shannon entropy 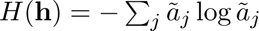 of the normalized absorbed spectrum; on the 17-signature curated panel Shannon entropy and IPR are rank-equivalent across the 8 TCGA cancers (median Spearman *ρ*(*H,* IPR) = +0.95; Supplementary Table S18), confirming that the choice between IPR and Shannon entropy as the spectral-diversity diagnostic does not change the interpretation. For the BRCA ExprPC200 null model, random 30-gene sets yield IPR near the expected-random anchor reported in Supplementary Table S4; the same random baseline yields *c* = 0.647 ± 0.110 (mean ± SD over 500 replicates), which we quote alongside the IPR values as a readability aid.

#### Link between spectral decomposition and the residual-ratio trajectory

When the null model is taken to be the leading *k* expression PCs (ExprPC*k*), the residual ratio is a simple partial sum of the absorbed spectrum:

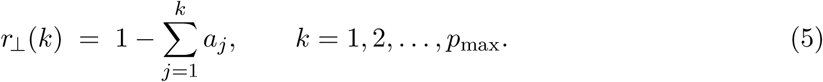

The sequence {*r*_⊥_(*k*)} is therefore a monotonically non-increasing, piecewise-linear function of the null-model dimensionality, and its rate of decay from *k* to *k* + 1 equals *a_k_*_+1_—the squared projection of **h** onto the (*k* + 1)-th expression PC. Two consequences follow directly. First, a signature whose direction is a linear combination of exactly *m* leading expression PCs (call it an *m-mode-aligned* signature) satisfies 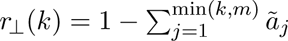, so *r*_⊥_ is flat for *k > m* and the trajectory shape identifies *m* as the effective absorbing rank. Second, a signature whose absorption is *diffuse* over many PCs has trajectory *r*_⊥_(*k*) ≈ 1 − *k/*IPR(**h**) to first order, so the slope of the trajectory at small *k* directly reports the signature’s IPR relative to the null-model size. These two regimes—sharply absorbing at small *k* versus slow, linear decay—are the operational content of the “trajectory is the load-bearing signal” claim in the main text: the trajectory shape, not any single *r*_⊥_(*k*), is the object that is computed, bootstrapped, and reported throughout this paper.

### 4.8 Cross-validated dimensionality selection

We ran a 5-fold cross-validation sweep over *k* ∈ {5, 10, 20, 50, 100, 200, 300} to select the null-model dimensionality. For each fold we fit PCA on the training split (80% of samples), projected the training basis onto the held-out split via the out-of-sample gene-space extension, re-orthonormalized via QR in the test sample-space, and reported two per-*k* statistics averaged across signatures and folds: (i) the training–test gap 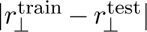 and (ii) the ranking stability *ρ* between training and test signature orderings. The CV-optimal *k*^∗^ was defined as the value minimising the train–test gap subject to a ranking-stability threshold of *ρ >* 0.90. The threshold *ρ >* 0.90 is a researcher-selected stability cutoff adopted *a priori* for its interpretability and is not pre-registered; a more conservative cutoff (e.g., *ρ >* 0.95) would tighten the CV-admissible range around *k* = 50 in several cancers, whereas a more permissive cutoff (e.g., *ρ >* 0.85) would widen it. We therefore do not treat the CV output as a formal selection rule; ExprPC50 is retained as the primary interpretive operating point across all cancers as a design choice (Methods §Null model hierarchy), and per-cancer CV *k*^∗^ values in Supplementary Table S12 are reported as supporting evidence rather than as a mandate. The procedure was applied independently in all 8 primary-analysis cancer types (BRCA, LUAD, COAD, OV, PRAD, SKCM, UCEC, HNSC); per-cancer values are reported in Supplementary Table S12. BRCA selects *k*^∗^ = 50 (gap 0.025, stability 0.98) as the strict CV optimum, matching the ExprPC50 operating point used throughout this work. LUAD, COAD, PRAD, UCEC, and HNSC all select *k*^∗^ = 20 (stability 0.92–0.96 at *k* = 20). SKCM selects *k*^∗^ = 5, and OV selects *k*^∗^ = 200 because its smaller sample size (*N* = 300) pushes the train–test gap minimum to the right-hand end of the grid. ExprPC50 is therefore the strict CV optimum for BRCA and lies inside the CV-admissible range (stability ≥ 0.87 at *k* = 50) for seven of the eight primary cancers. We use it as the primary interpretive operating point across all cancers and ExprPC200 as an upper-bound sensitivity level.

### 4.9 Formal significance testing

For the tier-level comparison, we used the Kruskal–Wallis test across the five validation tiers and report its rank-based *ε*^2^ effect size computed as *ε*^2^ = *H/*(*N* − 1) where *H* is the Kruskal– Wallis statistic and *N* is the total sample size. A complementary two-sided Mann–Whitney *U* test compared Tier 1 residual ratios against all other tiers combined, reported with the rank-biserial correlation *r* = 1 − 2*U/*(*n*_1_*n*_2_) as effect size. Pairwise Dunn’s post-hoc tests with Benjamini–Hochberg correction are reported in Supplementary Table S4.

For the permutation-based significance test, each signature’s residual ratio was compared against *n* = 500 size-matched random gene sets (or, for Reactome, *n* = 100 due to computational cost); the one-sided *p*-value is the fraction of random sets with residual ratio ≤ the signature’s ratio. All *p*-values were corrected for multiple testing using the Benjamini–Hochberg (BH) procedure [25]. We define three BH families consistent with how the tests are interpreted in the main text:

i. *pan-cancer curated family* — the 17 curated benchmark entries tested within each (cancer, null-model) slice (family size = 17 per slice, 8 cancer × 8 null-model = 64 independent slices); no cross-slice correction is applied. Used for Figure 4a and the “4–11 per cancer” count at ExprPC50;
ii. *Hallmark family* — all 50 MSigDB Hallmark gene sets tested within each (cancer, null-model) slice (family size = 50 per slice); no cross-slice correction is applied. Used for Hallmark counts in Results §Public pathway collections;
iii. *Reactome family* — all 1,181 Reactome 2022 pathways tested within each (cancer, null-model) slice (family size = 1,181 per slice); no cross-slice correction is applied. Used for Reactome counts.

Counts described in the Results as “significantly below the matched random baseline” refer to BH-adjusted permutation *p <* 0.05 within the relevant (cancer, null-model)-level family. Because Reactome uses 100 permutations against a 1,181-test BH family, the minimum resolvable BH-adjusted *p*-value is bounded below by the family size divided by the permutation count (≈ 0.012); we therefore interpret Reactome counts as descriptive indicators of pervasive absorption rather than fine-grained inferential statements. In the main text, counts of signatures falling below the matched random mean are reported as descriptive summaries, whereas Figure 4a and related inferential statements refer to permutation significance after BH correction. Both raw and adjusted *p*-values are reported where applicable.

### 4.10 Null model robustness analysis

To test sensitivity to null model specification, we constructed 12 alternative null models spanning three families: biological covariates (proliferation PC1 through PC20, immune PC10, combined proliferation + immune), and agnostic expression PCs (ExprPC20 through ExprPC300). We expanded the biological null family with four distinct covariate axes: proliferation (49 marker genes [85]), immune (29 markers), cell cycle (38 genes), and stromal (30 genes). Each axis was represented by its top 10 PCs. A combined all-biological null (40 PCs) was constructed by concatenating all four axes. Signature rankings were compared across all null model pairs using Spearman rank correlation.

### 4.11 Systematic overlap analysis

For each of the 16 curated pathway signatures and the housekeeping control, we computed the fraction of its genes present in any MSigDB Hallmark gene set (Hallmark coverage). We then removed all Hallmark-overlapping genes and recomputed residual ratios at the proliferation PC1 null model level, comparing original and adjusted ratios to quantify overlap-driven inflation. The correlation between Hallmark coverage and Δ*r*_⊥_ was assessed by Spearman rank correlation across all 17 benchmark entries.

### 4.12 Matched random baselines

To ensure that “below random baseline” claims are not artifacts of gene expression level or variance properties, we constructed three types of random baselines for each curated signature: (i) *size-matched*: *k* genes drawn uniformly from the gene universe; (ii) *expression-matched*: for each signature gene, a random gene was drawn from the 50 genes with most similar mean expression; (iii) *size+variance-matched*: jointly minimizing z-scored distance in mean expression and variance. Each baseline type was computed with 300 replicates per signature.

### 4.13 Statistical analysis

Bootstrap confidence intervals for cell-level residual ratios were computed using *B* = 1,000 sample-level resamples with replacement in a fixed-gene-space-basis procedure. The top-50 right-singular-vector basis **V**_50_ ∈ R*^G^*^×50^ was fit once on the full centered expression matrix (orthonormal in gene space by construction). For each bootstrap replicate, sample indices were resampled with replacement, and the resampled centered expression matrix was projected onto **V**_50_ to obtain per-sample scores, which were re-orthonormalized via QR decomposition to yield a valid orthonormal basis **Q***_b_* of the same 50-dimensional subspace in the resampled sample space. The signature direction **h***_b_* on the resampled samples was then projected onto **Q***_b_* to obtain *r*_⊥_. The 95% confidence interval was taken as the 2.5th and 97.5th percentiles of the 1,000-replicate distribution. This is a *fixed-basis* procedure: it captures sample-level uncertainty in the signature projection given a fixed gene-space subspace **V**_50_, but does not capture uncertainty in the subspace itself. A strictly more conservative alternative would be a *basis-refit* bootstrap in which the top-50 right-singular-vector basis is re-estimated on every resampled expression matrix; the basis-refit procedure would yield systematically wider cell-level CIs because it additionally propagates the variance of the SVD decomposition into each residual ratio. We chose the fixed-basis procedure because the gene-space basis **V**_50_ is what anchors the framework’s null-model definition, and because the trajectory-shape bootstrap (below) already saturates the stability question for the paper’s primary signal; cell-level CIs reported in Supplementary Table S5 should therefore be read as lower bounds on the true sampling uncertainty. Per-cell bootstrap 95% CIs for the Table 1 primary ExprPC50 residual ratios are reported in Supplementary Table S5. We additionally ran a *trajectory-shape bootstrap* for all 17 curated entries in BRCA: 200 sample-level resamples were drawn under the same procedure, and for each resample we computed the full trajectory *r*_⊥_(*k*) at *k* ∈ {1, 5, 10, 20, 50, 100, 200}. For every signature we then computed the mean pairwise Pearson correlation across the 200 trajectory-shape vectors as a measure of trajectory-shape stability; per-signature numbers are reported in Supplementary Table S14. Throughout the Results, *cross-cancer median* refers to the median over the 8 TCGA cancer types of the per-cancer point estimate, and *cross-cancer range* refers to the minimum and maximum per-cancer point estimates. For the METABRIC cross-cohort concordance, the 95% confidence interval for the Spearman rank correlation was computed via the Fisher *z*-transformation, 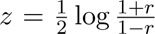, using 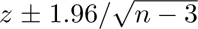 and back-transformed via tanh. For the gene swap test, we replaced signature genes with random genes in increments of 10% and repeated each swap level 30 times. Within-subtype analysis stratified BRCA samples by PAM50 molecular subtype using ExprPC20 as the null model. All random number generation used a fixed seed (42) for reproducibility, and the full reference implementation is available at the project repository listed in the Declarations.

## 5 Conclusions

The principal empirical finding is a consistent absolute gap in residual ratio between curated biological signatures and size-matched random 30-gene baselines at ExprPC50: across 8 TCGA cancers, the 17 curated pathway signatures have mean residual ratios 18–43% below the random baseline in every cancer (Supplementary Table S16). A secondary, descriptive observation is that, across the 17 curated signatures within each cancer, the ExprPC50 residual ratio is negatively correlated with the top-5 absorption concentration at median Spearman *ρ* = −0.71 (range −0.59 in PRAD to −0.89 in SKCM; all 8 cancers significant at *p <* 0.05). We report this correlation as a descriptive geometric property of the null-model coordinate system rather than as a biological law, because 1,000 random 30-gene draws per cancer reproduce the same pan-cancer median (*ρ* = −0.73; Supplementary Table S16), so any 30-gene direction inherits a similar correlation from the leading expression-PC axes. The correlation is robust to tumor-purity, immune, and stromal confounding: when the null basis is rebuilt on an immune-PC1 ⊕ stromal-PC1 ⊕ proliferation-PC1 anchor block plus the top-47 residual PCs, the per-cancer *ρ* becomes more negative rather than shallower (median −0.86; Supplementary Table S17), which rules out compositional nuisance as the driver. Signatures whose absorbed variance is concentrated on a small number of dominant expression-PC axes are the most strongly absorbed, regardless of validation tier; signatures whose absorption is distributed across many axes retain more residual variance. Under this reading, signature absorption is primarily a *geometric* property of how the signature direction aligns with the leading expression-PC axes of the data, and the co-occurrence of well-validated drivers (TP53) and independently validated cellular programs (autophagy, ferroptosis) in the same upper-residual-ratio band is a consequence of spectral geometry rather than an inconsistency. The corresponding trajectory shape *r*_⊥_(*k*) is empirically bootstrap-invariant across 200 sample-level resamples at mean pairwise Pearson correlation 0.999 (minimum 0.998, maximum 1.000 across the 17 entries); individual cell-level values at a single null-model dimensionality are also tight but not deterministic (BRCA median bootstrap 95% CI half-width 0.019, LUAD median 0.024 at *B* = 1,000 fixed-basis resamples, overall range ∼ 0.002– 0.053). The head-to-head reproduction of the Berglund uniqueness metric at three scales—17 curated entries, 50 Hallmark gene sets, and 1,361 Reactome pathways from the most recent Enrichr snapshot (versus the 1,181-pathway Reactome 2022 set used for the primary audit under the same 15–500 gene filter)—yields Spearman correlations in the range 0.39–0.67 and shows the two diagnostics capture closely related but non-identical aspects of the same absorbed-variance decomposition. A causal simulation further shows that residual ratio at any single null-model level cannot, on its own, adjudicate whether a non-zero residual reflects genuine confounder-independence or a confounded signal whose latent variables are only partially captured by the top principal components. Together, these observations motivate the framework’s practical output: the **trajectory shape** and the **curated-vs-random magnitude gap**, read jointly, are the load-bearing quantities—not the value of *r*_⊥_ at any single null-model dimensionality. We therefore propose residual-ratio auditing as a complementary audit layer to existing pathway-scoring and experimental-validation workflows, and stress that it does not replace clinical evaluation or experimental perturbation. A deterministic reference implementation, shipping both a pinned Python environment and an equivalent container image, is provided at the project repository (Declarations).

## Declarations

### Ethics approval and consent to participate

Not applicable. This study uses only previously published, fully de-identified public data obtained from the TCGA PanCancer Atlas and the METABRIC breast cancer study.

### Consent for publication

Not applicable.

### Availability of data and materials

All gene expression data supporting the conclusions of this article are openly available from cBioPortal (https://www.cbioportal.org). The primary analysis used the PanCancer Atlas RNA-Seq V2 RSEM studies for eight cancer types: breast (BRCA), lung adenocarcinoma (LUAD), colorectal (COAD), ovarian (OV), prostate (PRAD), skin melanoma (SKCM), uterine corpus endometrial (UCEC), and head and neck (HNSC). A supplementary ExprPC50 analysis additionally used the glioblastoma (GBM) and pancreatic (PAAD) PanCancer Atlas studies. External validation used the METABRIC breast cancer cohort from the same portal. All datasets were downloaded from cBioPortal between October 2025 and March 2026. The full list of PanCancer Atlas study identifiers, file names, and download timestamps is provided in Supplementary Table S15. Gene lists for all curated and control signatures, together with their provenance, are given in Supplementary Table S1.

A deterministic reference implementation of the residual-ratio framework, together with all scripts needed to regenerate the benchmark outputs and main text figures from the cBioPortal downloads above, is openly available at the project repository listed below. The implementation ships with a version-pinned Python 3.11 environment and an equivalent container image (Docker and Apptainer), so that all reported numerical values can be reproduced deterministically under a fixed random seed regardless of the host operating system.

Project name: Signature Audit.

Project home page: https://github.com/yudabi/signature-audit. Software release: v0.1.0.

Archived version: Zenodo DOI 10.5281/zenodo.pending (to be activated via the GitHub-Zenodo integration prior to final submission).

Operating systems: platform independent; tested on macOS and Linux and inside the container image.

Programming language: Python ≥ 3.11. License: MIT.

Restrictions to use by non-academics: none.

### Authors’ contributions

Y.B. conceived the study, developed the framework, performed all analyses, and wrote the manuscript. Y.Z. and C.Z. contributed to data curation and analysis.

V.D.C. supervised the project and revised the manuscript. All authors read and approved the final manuscript.

## Acknowledgements

The authors acknowledge the TCGA Research Network and the METABRIC Consortium for making the underlying expression data publicly available through cBioPortal, and thank the developers of sigQC, variancePartition, GSVA, singscore, and PROGENy for open-source implementations that made benchmarking straightforward.

## Supplementary Material

**Table S1:**
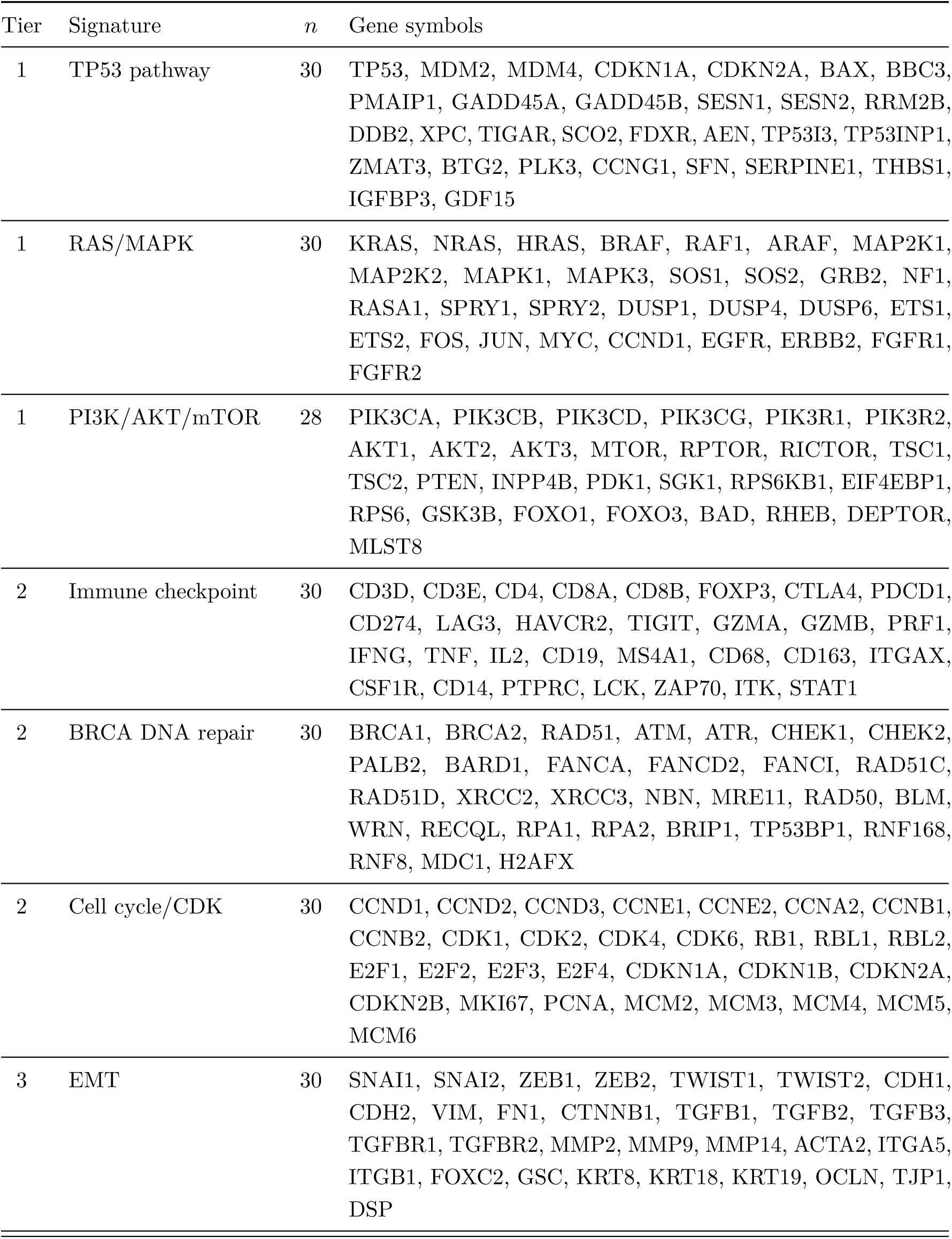
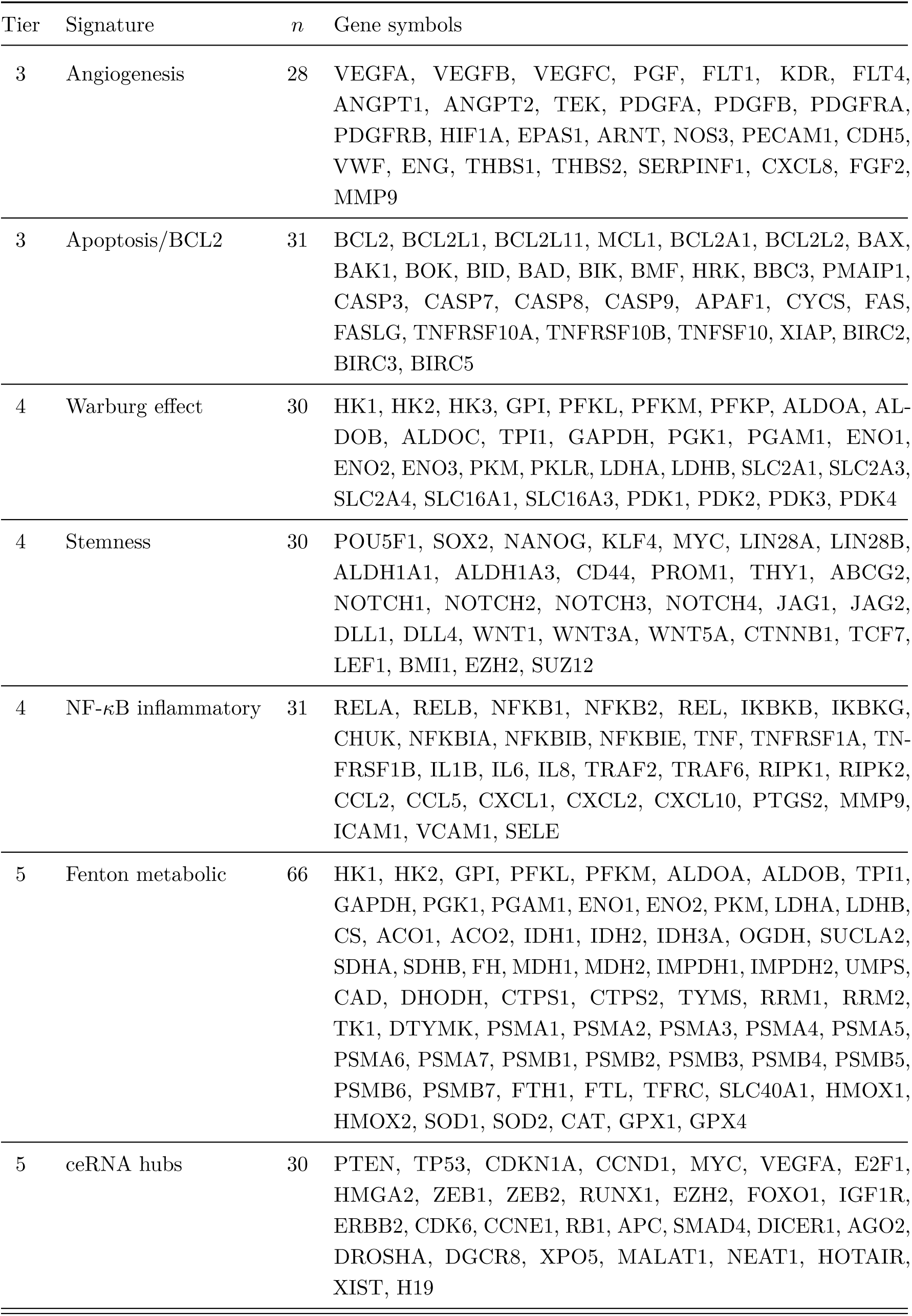
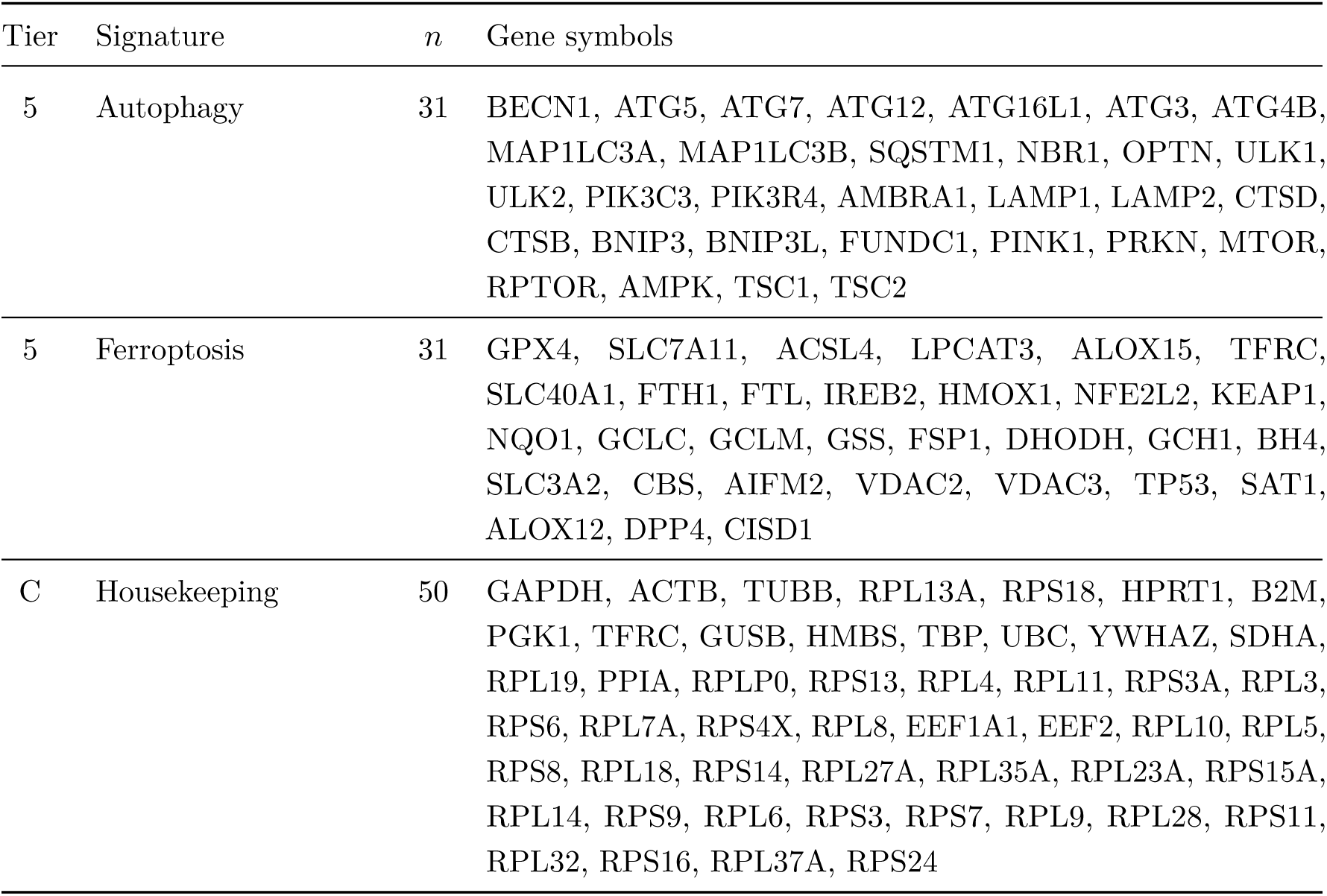
Curated gene signature gene lists. Explicit gene lists for all 16 curated pathway signatures and the housekeeping control used in the residual decomposition analysis.

**Table S2:** Null model covariate gene lists. Gene lists used to construct the biological null model covariates.

| Covariate axis | $n$ | Gene symbols |
| --- | --- | --- |
| Proliferation markers | 50 | ANLN, AURKA, AURKB, BIRC5, BUB1, BUB1B, CCNA2, CCNB1, CCNB2, CDC20, CDC6, CDCA5, CDCA8, CDK1, CENPA, CEP55, DLGAP5, ECT2, FOXM1, GINS2, GMNN, HJURP, KIF11, KIF20A, KIF2C, KIF4A, MCM2, MCM3, MCM4, MCM5, MCM6, MELK, MKI67, NDC80, NUF2, NUSAP1, ORC6, PCNA, PLK1, POLA2, POLD1, PRC1, RFC4, RRM2, TK1, TOP2A, TPX2, TTK, TYMS, UBE2C |
| Immune markers | 30 | CD14, CD163, CD19, CD274, CD3D, CD3E, CD4, CD68, CD8A, CD8B, CSF1R, CTLA4, FOXP3, GZMA, GZMB, HAVCR2, IFNG, IL2, ITGAX, ITK, LAG3, LCK, MS4A1, PDCD1, PRF1, PTPRC, STAT1, TIGIT, TNF, ZAP70 |
| Cell cycle | 38 | CDK1, CDK2, CDK4, CDK6, CCNA2, CCNB1, CCNB2, CCND1, CCNE1, CCNE2, CDC20, CDC25A, CDC25B, CDC25C, CDC45, E2F1, E2F2, E2F3, MCM2, MCM3, MCM4, MCM5, MCM6, MCM7, AURKA, AURKB, PLK1, BUB1, BUB1B, MAD2L1, CHEK1, CHEK2, RB1, TP53, CDKN1A, CDKN2A, MKI67, PCNA |
| Stromal | 30 | FAP, PDGFRA, PDGFRB, COL1A1, COL1A2, COL3A1, FN1, ACTA2, VIM, THY1, DCN, LUM, SPARC, MMP2, MMP3, MMP9, TGFB1, TGFB2, TGFB3, LOX, LOXL2, POSTN, SNAI1, SNAI2, TWIST1, ZEB1, ZEB2, CDH2, S100A4, TAGLN |

**Table S3:** Residual ratios at the upper-bound ExprPC200 sensitivity level across 8 TCGA cancer types. Bold values indicate the highest residual ratio per cancer type. The “Random” row shows the mean ratio of 200 random 100-gene sets.

| Tier | Signature | BRCA | LUAD | COAD | OV | PRAD | SKCM | UCEC | HNSC |
| --- | --- | --- | --- | --- | --- | --- | --- | --- | --- |
| 1 | TP53 pathway | 0.140 | **0.099** | 0.078 | 0.043 | 0.044 | **0.096** | 0.073 | **0.110** |
| 1 | RAS/MAPK | 0.073 | 0.063 | 0.040 | 0.026 | 0.020 | 0.074 | 0.044 | 0.047 |
| 1 | PI3K/AKT/mTOR | 0.085 | 0.052 | 0.040 | 0.041 | 0.038 | 0.055 | 0.060 | 0.054 |
| 2 | Immune checkpoint | 0.006 | 0.007 | 0.006 | 0.003 | 0.005 | 0.003 | 0.007 | 0.005 |
| 2 | BRCA DNA repair | 0.024 | 0.012 | 0.017 | 0.016 | 0.018 | 0.012 | 0.014 | 0.015 |
| 2 | Cell cycle/CDK | 0.025 | 0.024 | 0.052 | 0.021 | 0.018 | 0.053 | 0.026 | 0.046 |
| 3 | EMT | 0.032 | 0.024 | 0.010 | 0.009 | 0.014 | 0.043 | 0.038 | 0.019 |
| 3 | Angiogenesis | 0.020 | 0.016 | 0.008 | 0.009 | 0.010 | 0.017 | 0.023 | 0.013 |
| 3 | Apoptosis/BCL2 | 0.080 | 0.045 | 0.026 | 0.023 | 0.041 | 0.041 | 0.047 | 0.055 |
| 4 | Warburg effect | 0.085 | 0.052 | 0.078 | 0.037 | 0.046 | 0.053 | 0.074 | 0.054 |
| 4 | Stemness | 0.066 | 0.048 | 0.060 | 0.061 | 0.026 | 0.061 | 0.081 | 0.043 |
| 4 | NF-κB inflammatory | 0.031 | 0.032 | 0.021 | 0.011 | 0.017 | 0.019 | 0.030 | 0.026 |
| 5 | Fenton metabolic | 0.044 | 0.027 | 0.036 | 0.013 | 0.031 | 0.030 | 0.025 | 0.035 |
| 5 | ceRNA hubs | 0.115 | 0.048 | 0.046 | 0.034 | 0.055 | 0.044 | 0.064 | 0.081 |
| 5 | Autophagy | **0.168** | 0.097 | 0.068 | **0.062** | **0.064** | 0.072 | **0.084** | 0.103 |
| 5 | Ferroptosis | 0.161 | 0.064 | **0.117** | 0.058 | 0.062 | 0.074 | 0.079 | 0.051 |
| C | Housekeeping | 0.042 | 0.033 | 0.033 | 0.015 | 0.019 | 0.031 | 0.040 | 0.023 |
| C | Random ($n = 200$) | 0.077 | 0.050 | 0.036 | 0.039 | 0.032 | 0.038 | 0.050 | 0.036 |

**Table S4:** Pairwise Dunn’s post-hoc tests with Benjamini–Hochberg correction for the tier gradient at ExprPC50. Tier ratios pooled across all 8 TCGA cancer types (*n* = 24 each for Tiers 1–4; *n* = 32 for Tier 5). The omnibus Kruskal–Wallis test reports *H* = 63.7, df = 4, *ε*^2^ = 0.50, *p* = 4.9 10−13. Tier 1 is significantly shifted above Tiers 2 and 3 but not separable from Tiers 4 and 5, reflecting a tier-wide distributional shift rather than a strict pairwise ordering. Tier 5 contains computationally assembled gene lists whose *underlying biological processes* (autophagy, ferroptosis) are independently well validated; the high Tier 5 residual ratios in particular are driven by the Autophagy and Ferroptosis gene lists rather than by the ceRNA hubs or Fenton metabolic lists.

| Tier pair | $p_{BH}$ | Significant at $\alpha = 0.05$? | Median (lower tier) | Median (higher tier) |
| --- | --- | --- | --- | --- |
| T1 vs T2 | $5.8 \times 10^{-8}$ | *** | 0.059 (T2) | 0.188 (T1) |
| T1 vs T3 | $1.6 \times 10^{-6}$ | *** | 0.072 (T3) | 0.188 (T1) |
| T1 vs T4 | 0.218 | ns | 0.157 (T4) | 0.188 (T1) |
| T1 vs T5 | 0.978 | ns | 0.188 (T1) | 0.202 (T5) |
| T2 vs T3 | 0.521 | ns | 0.059 (T2) | 0.072 (T3) |
| T2 vs T4 | $2.8 \times 10^{-5}$ | *** | 0.059 (T2) | 0.157 (T4) |
| T2 vs T5 | $1.3 \times 10^{-8}$ | *** | 0.059 (T2) | 0.202 (T5) |
| T3 vs T4 | $4.9 \times 10^{-4}$ | *** | 0.072 (T3) | 0.157 (T4) |
| T3 vs T5 | $3.9 \times 10^{-7}$ | *** | 0.072 (T3) | 0.202 (T5) |
| T4 vs T5 | 0.218 | ns | 0.157 (T4) | 0.202 (T5) |

Significance codes: \*\*\* $p < 0.001$ ; ns $p \geq 0.05$ . Mann–Whitney $U$ test Tier 1 vs rest combined: $U = 1830$ , rank-biserial $r = 0.47$ , $p = 3.9 \times 10^{-4}$ .

**Table S5:** Sample-only bootstrap 95% confidence intervals for Table 1 residual ratios at ExprPC50. For each cell, we fit the top-50 PCA basis once on the full centered log_2_(RSEM+1) expression matrix, precomputed the unit signature direction **h** in sample space, and then ran 200 sample-level bootstrap replicates in which sample indices were resampled with replacement and the residual ratio recomputed against the restricted PCA scores. The point estimate is the residual ratio on full data; columns 3–4 report the 2.5th and 97.5th percentiles of the 200-replicate distribution and column 5 the 95% CI half-width. All values are reproducible from the bootstrap routine archived in the project repository.

| Cancer | Signature | $r_{\perp}$ (point) | 2.5% | 97.5% | Half-width |
| --- | --- | --- | --- | --- | --- |
| BRCA | TP53 pathway | 0.306 | 0.263 | 0.325 | 0.031 |
| BRCA | RAS/MAPK | 0.196 | 0.166 | 0.208 | 0.021 |
| BRCA | PI3K/AKT/mTOR | 0.185 | 0.155 | 0.197 | 0.021 |
| BRCA | Immune checkpoint | 0.017 | 0.014 | 0.021 | 0.003 |
| BRCA | BRCA DNA repair | 0.057 | 0.049 | 0.062 | 0.007 |
| BRCA | Cell cycle/CDK | 0.069 | 0.060 | 0.073 | 0.006 |
| BRCA | EMT | 0.065 | 0.054 | 0.070 | 0.008 |
| BRCA | Angiogenesis | 0.050 | 0.041 | 0.054 | 0.007 |
| BRCA | Apoptosis/BCL2 | 0.188 | 0.157 | 0.203 | 0.023 |
| BRCA | Warburg effect | 0.251 | 0.214 | 0.259 | 0.022 |
| BRCA | Stemness | 0.123 | 0.103 | 0.133 | 0.015 |
| BRCA | NF-κB | 0.104 | 0.089 | 0.110 | 0.010 |
| BRCA | Fenton metabolic | 0.121 | 0.102 | 0.129 | 0.014 |
| BRCA | ceRNA hubs | 0.245 | 0.206 | 0.262 | 0.028 |
| BRCA | Autophagy | 0.410 | 0.354 | 0.429 | 0.038 |
| BRCA | Ferroptosis | 0.321 | 0.272 | 0.339 | 0.034 |
| BRCA | Housekeeping | 0.171 | 0.146 | 0.179 | 0.017 |
| LUAD | TP53 pathway | 0.370 | 0.277 | 0.387 | 0.055 |
| LUAD | RAS/MAPK | 0.237 | 0.182 | 0.248 | 0.033 |
| LUAD | PI3K/AKT/mTOR | 0.192 | 0.143 | 0.201 | 0.029 |
| LUAD | Immune checkpoint | 0.022 | 0.017 | 0.030 | 0.006 |
| LUAD | BRCA DNA repair | 0.047 | 0.036 | 0.053 | 0.008 |
| LUAD | Cell cycle/CDK | 0.081 | 0.061 | 0.086 | 0.013 |
| LUAD | EMT | 0.065 | 0.049 | 0.072 | 0.011 |
| LUAD | Angiogenesis | 0.074 | 0.058 | 0.080 | 0.011 |
| LUAD | Apoptosis/BCL2 | 0.120 | 0.090 | 0.129 | 0.019 |
| LUAD | Warburg effect | 0.232 | 0.178 | 0.238 | 0.030 |
| LUAD | Stemness | 0.127 | 0.095 | 0.140 | 0.023 |
| LUAD | NF-κB | 0.158 | 0.117 | 0.168 | 0.026 |
| LUAD | Fenton metabolic | 0.106 | 0.079 | 0.116 | 0.019 |
| LUAD | ceRNA hubs | 0.153 | 0.116 | 0.162 | 0.023 |
| LUAD | Autophagy | 0.315 | 0.230 | 0.326 | 0.048 |
| LUAD | Ferroptosis | 0.183 | 0.135 | 0.190 | 0.028 |
| LUAD | Housekeeping | 0.140 | 0.104 | 0.158 | 0.027 |

**Table S6:** Causal DAG sensitivity analysis — 27 parameter configurations. One-at-a-time sweep around the base configuration (*N* = 500, *G* = 5,000, *k*_sig_ = 30, confounder strength = 0.8, propagation strength = 1.5, noise scale = 1.0, *n*_downstream_ = 500). Each configuration was run with 20 independent Monte-Carlo replicates. The gap is the mean confounder-collinear residual ratio minus the mean mediator-driven residual ratio at ExprPC5. “Dist.” = distinguishable (gap positive and 95% CI not containing zero). Across all 27 configurations, 24 (89%, Clopper–Pearson 95% CI 0.71–0.98) preserve distinguishability at ExprPC5. (mean ± SD *Failures:* 3 of 27 configurations fail to preserve distinguishability (confounder strength ≥ 2.0 dominates both scenarios equally; noise scale 0.3 produces near-zero signal in both scenarios). Entries marked “Yes^∗^” have negative gap (mediator-driven above confounder-collinear) but remain distinguishable; the direction reversal occurs because weak propagation strength allows the pathway to drive downstream expression more strongly than the confounder in the low-*k* regime. For full reproducibility, the exact parameter values, seeds, and per-replicate results are available in the project repository.

| Swept parameter | Value | Gap @ PC5<br>(mean ± SD) | Gap @ PC200<br>(mean ± SD) | Dist. |
| --- | --- | --- | --- | --- |
| Confounder strength | 0.3 | $0.612 \pm 0.141$ | $0.295 \pm 0.056$ | Yes |
| Confounder strength | 0.5 | $0.398 \pm 0.140$ | $0.188 \pm 0.050$ | Yes |
| Confounder strength | 0.8 (base) | $0.201 \pm 0.104$ | $0.094 \pm 0.040$ | Yes |
| Confounder strength | 1.2 | $0.076 \pm 0.068$ | $0.042 \pm 0.028$ | Yes |
| Confounder strength | 2.0 | $-0.011 \pm 0.039$ | $-0.003 \pm 0.016$ | No |
| Confounder strength | 3.0 | $-0.044 \pm 0.027$ | $-0.014 \pm 0.011$ | No |
| Propagation strength | 0.3 | $-0.553 \pm 0.182$ | $-0.168 \pm 0.075$ | Yes* |
| Propagation strength | 0.6 | $-0.167 \pm 0.235$ | $-0.051 \pm 0.086$ | Yes* |
| Propagation strength | 1.0 | $0.127 \pm 0.117$ | $0.055 \pm 0.042$ | Yes |
| Propagation strength | 1.5 (base) | $0.201 \pm 0.104$ | $0.094 \pm 0.040$ | Yes |
| Propagation strength | 2.0 | $0.230 \pm 0.098$ | $0.109 \pm 0.039$ | Yes |
| Propagation strength | 3.0 | $0.252 \pm 0.094$ | $0.120 \pm 0.039$ | Yes |
| Noise scale | 0.3 | $0.029 \pm 0.016$ | $0.014 \pm 0.008$ | No |
| Noise scale | 0.5 | $0.072 \pm 0.039$ | $0.034 \pm 0.017$ | Yes |
| Noise scale | 1.0 (base) | $0.201 \pm 0.104$ | $0.094 \pm 0.040$ | Yes |
| Noise scale | 1.5 | $0.293 \pm 0.150$ | $0.138 \pm 0.058$ | Yes |
| Noise scale | 2.0 | $0.333 \pm 0.173$ | $0.157 \pm 0.066$ | Yes |
| Pathway gene-set size $k_{\text{sig}}$ | 10 | $0.186 \pm 0.063$ | $0.090 \pm 0.034$ | Yes |
| Pathway gene-set size $k_{\text{sig}}$ | 20 | $0.163 \pm 0.070$ | $0.080 \pm 0.030$ | Yes |
| Pathway gene-set size $k_{\text{sig}}$ | 30 (base) | $0.201 \pm 0.104$ | $0.094 \pm 0.040$ | Yes |
| Pathway gene-set size $k_{\text{sig}}$ | 50 | $0.121 \pm 0.125$ | $0.057 \pm 0.049$ | Yes |
| Pathway gene-set size $k_{\text{sig}}$ | 100 | $0.154 \pm 0.132$ | $0.074 \pm 0.051$ | Yes |
| Num downstream $n_{\text{downstream}}$ | 50 | $0.203 \pm 0.102$ | $0.095 \pm 0.040$ | Yes |
| Num downstream $n_{\text{downstream}}$ | 100 | $0.202 \pm 0.104$ | $0.094 \pm 0.040$ | Yes |
| Num downstream $n_{\text{downstream}}$ | 200 | $0.201 \pm 0.104$ | $0.094 \pm 0.040$ | Yes |
| Num downstream $n_{\text{downstream}}$ | 500 (base) | $0.201 \pm 0.104$ | $0.094 \pm 0.040$ | Yes |
| Num downstream $n_{\text{downstream}}$ | 1000 | $0.201 \pm 0.104$ | $0.094 \pm 0.040$ | Yes |

**Table S7:**
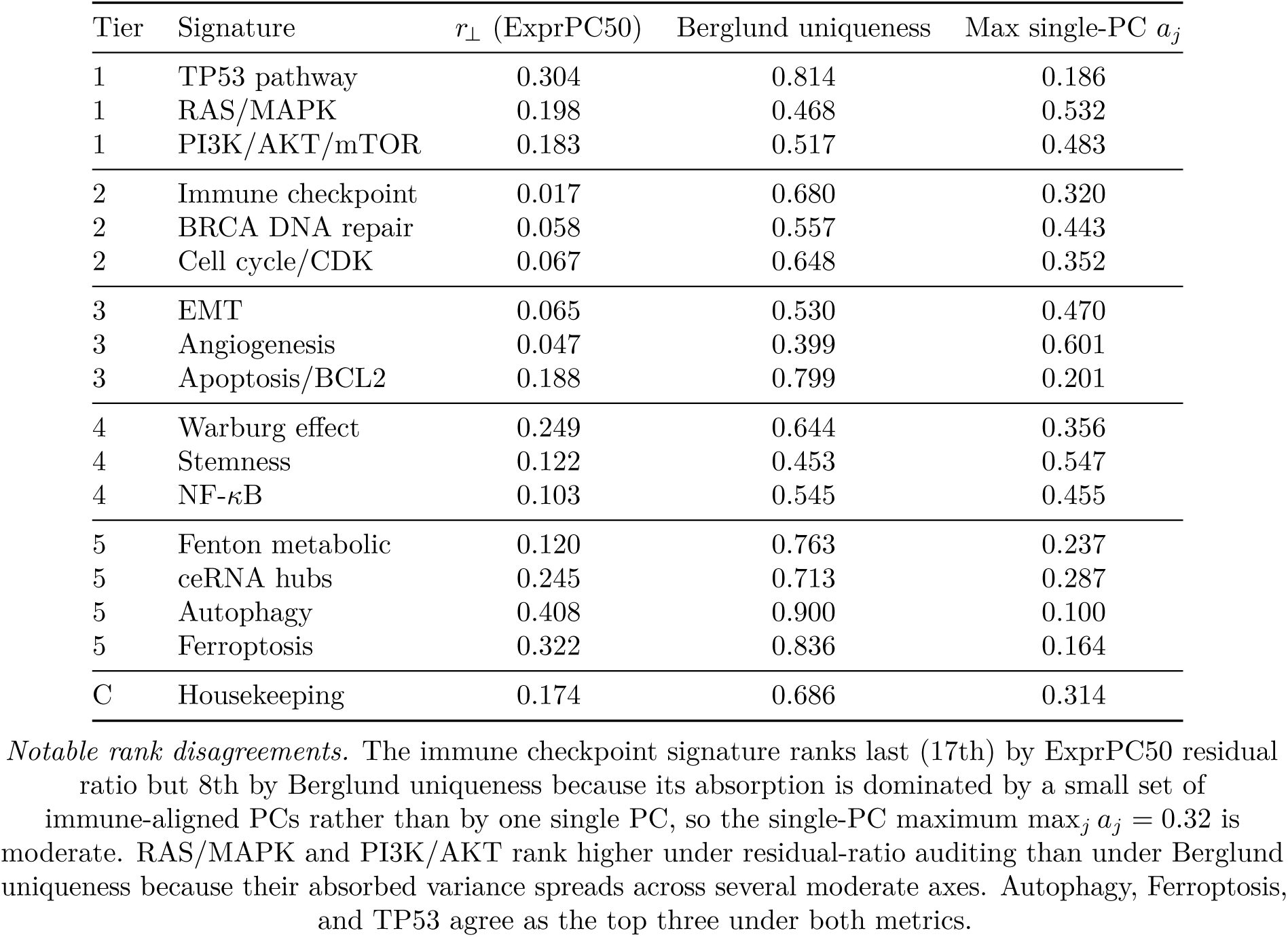
Berglund-style uniqueness versus ExprPC50 residual ratio for the 17-entry curated benchmark in BRCA. Berglund-style uniqueness is operationalized as 1 max*_j_ a_j_*where *a_j_* is the squared projection of the unit signature direction onto the *j*-th leading expression PC (high uniqueness = no single global PC captures a large share of the signature). Spearman rank correlation between the two metrics: *ρ* = 0.57, 95% Fisher-*z* CI 0.12–0.82, *p* = 0.018, *n* = 17.

**Table S8:** ExprPC50 residual ratios for GBM and PAAD (supplementary analysis, cancer types with *N <* 200 samples). These cancer types were excluded from the primary analysis because the ExprPC200 upper-bound null model requires *N* ≥ 201 + 2 samples (see Methods § Data sources). At the primary ExprPC50 operating point they are tractable and show the same qualitative tier gradient as the 8 primary cancers: Tier 1 TP53 and Tier 5 Autophagy/Ferroptosis occupy the top positions in both. This supplementary table therefore confirms that the primary conclusions are not altered by the exclusion.

| Tier | Signature | GBM ($N = 160$) | PAAD ($N = 177$) |
| --- | --- | --- | --- |
| 1 | TP53 pathway | 0.150 | 0.171 |
| 1 | RAS/MAPK | 0.089 | 0.084 |
| 1 | PI3K/AKT/mTOR | 0.106 | 0.043 |
| 2 | Immune checkpoint | 0.038 | 0.014 |
| 2 | BRCA DNA repair | 0.036 | 0.063 |
| 2 | Cell cycle/CDK | 0.042 | 0.059 |
| 3 | EMT | 0.055 | 0.020 |
| 3 | Angiogenesis | 0.048 | 0.030 |
| 3 | Apoptosis/BCL2 | 0.068 | 0.032 |
| 4 | Warburg effect | 0.103 | 0.113 |
| 4 | Stemness | 0.155 | 0.062 |
| 4 | NF-κB | 0.055 | 0.084 |
| 5 | Fenton metabolic | 0.057 | 0.047 |
| 5 | ceRNA hubs | 0.066 | 0.058 |
| 5 | Autophagy | **0.098** | **0.198** |
| 5 | Ferroptosis | 0.079 | **0.230** |
| C | Housekeeping | 0.044 | 0.081 |
| C | Random ($n = 200$) | 0.077 | 0.058 |

**Table S9:** Cumulative variance explained by the ExprPC50 and ExprPC200 null-model subspaces. Values are the fraction of total transcriptomic variance captured by the top-*k* principal components of the log_2_(RSEM+1)-transformed, median-filtered expression matrix for each cancer type (*G* ≈ 16,400–16,900 genes per cancer after median-filter). All values are reproducible from the per-cancer PCA pipeline archived in the project repository.

| Cancer | $N$ (samples) | Variance at $k = 50$ | Variance at $k = 200$ |
| --- | --- | --- | --- |
| BRCA | 1,082 | 61.6% | 78.9% |
| LUAD | 510 | 64.5% | 85.6% |
| COAD | 592 | 65.9% | 84.8% |
| OV | 300 | 63.5% | 92.1% |
| PRAD | 493 | 69.8% | 88.3% |
| SKCM | 443 | 66.3% | 87.4% |
| UCEC | 527 | 64.5% | 85.0% |
| HNSC | 515 | 68.7% | 87.1% |

*Cross-cancer averages: 64.6% at $k = 50$ , 85.9% at $k = 200$ . Both operating points capture the majority of transcriptomic variance; ExprPC50 is used as the primary operating point because it retains stable rankings across null-model choices, and ExprPC200 as an upper-bound sensitivity level. OV has the highest variance-at-200 (92.1%) because its smaller sample size ( $N = 300$ ) makes the top 200 PCs closer to the full-rank basis. All per-cancer PCA runs used the same pipeline ( $\log_2(\text{RSEM}+1)$ , median-filter at 1.0, randomized SVD, random\_state=42).*

**Table S10:**
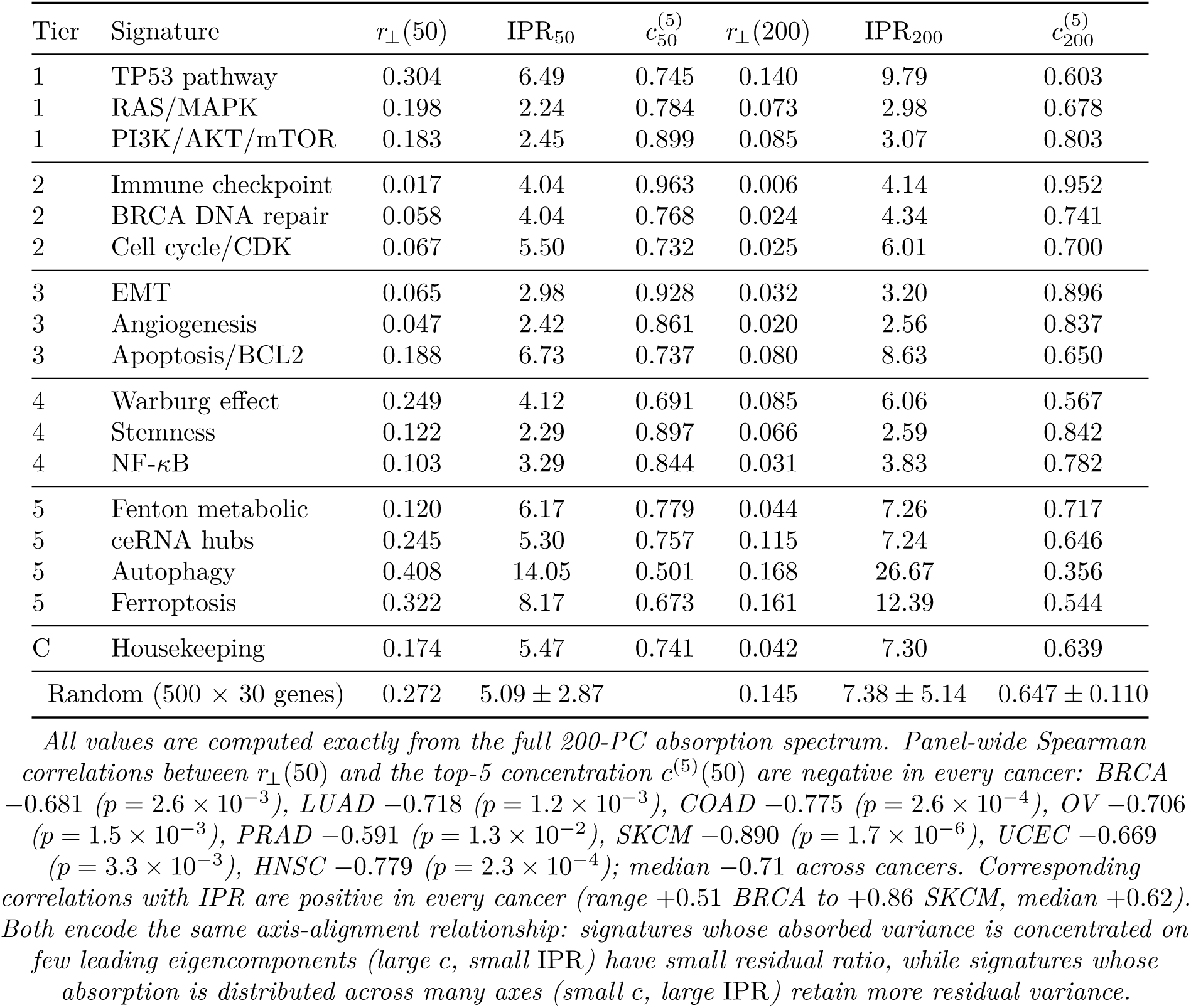
Panel-wide absorption concentration: IPR and top-5 concentration for all 17 curated benchmark entries. For each signature, we report the effective number of absorbing eigencomponents (inverse participation ratio, 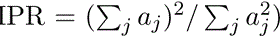, the top-5 concentration *c*, and the single-leading-axis contribution 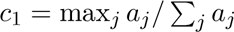, computed at the primary ExprPC50 operating point and at the upper-bound ExprPC200 level in BRCA. All statistics are derived from the full 200-PC per-signature absorption spectrum 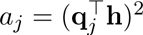 using the signature_audit reference implementation. The random-baseline row gives the mean ± SD of 500 size-matched 30-gene random sets in BRCA under the same ExprPC200 null model. The Spearman rank correlations at the foot of the table quantify the panel-wide relationship between concentration and residual ratio (see Results §Interpretive boundaries).

**Table S11:** Full-scale head-to-head between residual-ratio auditing and Berglund uniqueness. For each of the 17 curated benchmark entries, all 50 MSigDB Hallmark gene sets, and all 1,181 Reactome 2022 pathways (filtered to 15–500 genes), we computed the ExprPC50 residual ratio and Berglund’s uniqueness metric 1 max*_j_ a_j_* from the same top-200 expression-PC basis in BRCA and LUAD. Spearman rank correlation between the two metrics is reported at each scale with a Fisher-*z*-transformed 95% CI. At the *n* = 17 scale, the CI reproduces the prior report (*ρ* = 0.57, CI 0.12–0.82); at the *n* = 50 and *n* = 1,181 scales the CI tightens substantially, allowing a more definitive statement about the relationship between the two metrics.

| Scale | Cancer | $n$ | Spearman $\rho$ | Fisher-$z$ 95% CI | $p$-value |
| --- | --- | --- | --- | --- | --- |
| Curated 17-entry | BRCA | 17 | +0.57 | +0.12 – +0.82 | $1.8 \times 10^{-2}$ |
| Curated 17-entry | LUAD | 17 | +0.62 | +0.19 – +0.85 | $8.6 \times 10^{-3}$ |
| MSigDB Hallmark | BRCA | 50 | +0.39 | +0.12 – +0.60 | $5.2 \times 10^{-3}$ |
| MSigDB Hallmark | LUAD | 50 | +0.61 | +0.40 – +0.76 | $2.5 \times 10^{-6}$ |
| Reactome (filtered 15–500 genes) | BRCA | 1,361 | +0.55 | +0.51 – +0.59 | $< 10^{-100}$ |
| Reactome (filtered 15–500 genes) | LUAD | 1,361 | +0.67 | +0.64 – +0.70 | $< 10^{-100}$ |

*All numerical values are reproduced deterministically under the fixed random seed reported in the Methods, against the current Enrichr MSigDB Hallmark 2020 and Reactome 2024 libraries. The filtered Reactome library contains 1,361 pathways after the 15–500 gene-count filter, a small library update relative to the 1,181-pathway Reactome 2022 set used for the primary audit. The single qualitative disagreement observable across all three scales is the immune checkpoint gene set family: under residual-ratio auditing at ExprPC50 these sets rank near the bottom of the benchmark (strongly absorbed across the full 50-dimensional null-model subspace), while under Berglund uniqueness they rank only moderately low because $1 - \max_j a_j$ is insensitive to absorption distributed across two or three near-top immune-aligned PCs. Because the Hallmark and Reactome Fisher- $z$ CIs exclude zero by many orders of magnitude, the two metrics are neither perfectly redundant nor independent but capture closely related aspects of the same absorbed-variance decomposition.*

**Table S12:** Per-cancer 5-fold cross-validation sweep: train–test gap and ranking stability as functions of null-model dimensionality. *k*. For each of the 8 primary-analysis cancer types, we ran a 5-fold CV sweep over *k* 5, 10, 20, 50, 100, 200, 300, with training PCs computed by SVD of the training split and test samples projected onto the training gene-space basis and re-orthonormalized via QR before computing the residual ratio. Each row shows the mean training–test [jnline] across signatures and folds, and the mean Spearman ranking stability *ρ* between training and test signature orderings. The “CV-optimal *k*^∗^” column minimises the train–test gap subject to the stability constraint *ρ >* 0.90. BRCA selects *k*^∗^ = 50, matching the primary interpretive operating point used throughout this work; five of the remaining seven cancers select *k*^∗^ = 20; SKCM selects *k*^∗^ = 5 and OV selects *k*^∗^ = 200.

| Cancer | CV-optimal | | Train–test gap at $k =$ | | | Stability at $k =$ | |
| --- | --- | --- | --- | --- | --- | --- | --- |
| | $k^*$ | gap@$k^*$ | 5 | 50 | 200 | 5 | 50 |
| BRCA | 50 | 0.025 | 0.038 | 0.025 | 0.059 | 0.96 | 0.98 |
| LUAD | 20 | 0.045 | 0.061 | 0.054 | 0.035 | 0.93 | 0.92 |
| COAD | 20 | 0.036 | 0.057 | 0.043 | 0.036 | 0.93 | 0.95 |
| OV | 200 | 0.013 | 0.064 | 0.113 | 0.013 | 0.86 | 0.71 |
| PRAD | 20 | 0.034 | 0.051 | 0.038 | 0.024 | 0.95 | 0.90 |
| SKCM | 5 | 0.056 | 0.056 | 0.075 | 0.035 | 0.95 | 0.87 |
| UCEC | 20 | 0.046 | 0.059 | 0.054 | 0.038 | 0.92 | 0.87 |
| HNSC | 20 | 0.047 | 0.055 | 0.054 | 0.037 | 0.96 | 0.89 |

*All values are reproduced deterministically under the fixed random seed reported in the Methods. BRCA selects $k^* = 50$ as the strict CV optimum, matching the ExprPC50 operating point adopted throughout this work; the remaining six primary cancers (LUAD, COAD, PRAD, SKCM, UCEC, HNSC) have their CV-optimal $k^*$ in $\{5, 20\}$ , and ExprPC50 lies inside the CV-admissible range (ranking stability $\geq 0.87$ ) for all seven. OV is the only outlier, with $k^* = 200$ because its smaller sample size ( $N = 300$ ) pushes the train–test gap minimum to the right-hand end of the grid while ranking stability drops sharply for $k \geq 100$ .*

**Table S13:** Causal DAG simulation at *n*_rep_ = 100: tighter base-configuration CIs and the 27-configuration sensitivity sweep. We re-ran the DAG simulation of Methods § 4.6 on the base configuration with *n*_rep_ = 100 Monte-Carlo replicates (raised from 20 in the prior version of this work) using the same generative model. The earlier single-point claim “*r*_⊥_ 0.25 at ExprPC50” is preserved: the tighter-CI estimate is 0.233 with a 95% CI on the mean of [0.210, 0.256]. The 27-configuration sensitivity sweep is retained from the prior version as a one-at-a-time design across confounder strength, propagation strength, noise scale, gene-set size, and downstream gene count; its reported outcome is the binary “gap positive at ExprPC5” flag, which does not change materially at higher *n*_rep_.

| Scenario | $k$ | Prior ($n_{\text{rep}} = 20$) | New ($n_{\text{rep}} = 100$) | New 95% CI |
| --- | --- | --- | --- | --- |
| Confounder-collinear | 5 | $0.290 \pm 0.111$ | $0.273 \pm 0.136$ | $[0.246, 0.300]$ |
| Confounder-collinear | 50 | $0.251 \pm 0.099$ | $0.233 \pm 0.115$ | $[0.210, 0.256]$ |
| Confounder-collinear | 200 | $0.140 \pm 0.055$ | $0.074 \pm 0.051$ | $[0.064, 0.084]$ |
| Mediator-driven | 5 | $0.111 \pm 0.055$ | $0.128 \pm 0.100$ | $[0.108, 0.148]$ |
| Mediator-driven | 50 | $0.095 \pm 0.048$ | $0.110 \pm 0.086$ | $[0.093, 0.127]$ |
| Gap (conf – med) at ExprPC5 | | $\approx 0.18$ | 0.144 | $[0.111, 0.178]$ |
| 27-configuration sensitivity sweep: gap positive at ExprPC5 in 24/27 configurations (89%; Clopper–Pearson 95% CI $[0.71, 0.98]$; prior-version one-at-a-time design). | | | | |

*All base-configuration values in the middle two columns and the 95% CI column are reproduced deterministically under the fixed random seed reported in the Methods. The new estimates are close to the prior point estimates but carry tighter CIs by a factor of $\sqrt{100/20} \approx 2.2$ . The confounder $r_{\perp}$ (ExprPC200) values differ more noticeably between versions (0.140 vs 0.074) because at $k = 200$ the absorbed spectrum is already $\sim 93\%$ saturated and the remaining orthogonal variance is dominated by finite- $N$ noise, so Monte-Carlo noise at $n_{\text{rep}} = 20$ was the dominant source of the prior point estimate; the $n_{\text{rep}} = 100$ rerun places a lower but tighter value on the same quantity.*

**Table S14:** Trajectory-shape bootstrap stability for the 17 curated benchmark entries in BRCA. For each signature we ran 200 sample-level fixed-gene-space-basis bootstrap resamples, and for each resample computed the full trajectory *r*_⊥_(*k*) at *k* 1, 5, 10, 20, 50, 100, 200 . Columns 3–5 report the bootstrap standard deviation at *k* = 5, *k* = 50, and *k* = 200; column 6 reports the mean pairwise Pearson correlation across the 200 trajectory-shape vectors per signature. Values near 1.0 indicate that the trajectory shape is effectively deterministic at the sample level. Across all 17 entries the mean pairwise shape correlation has mean 0.9991, median 0.9992, minimum 0.9982, maximum 1.0000—the trajectory shape is empirically bootstrap-invariant.

| Tier | Signature | $\sigma_{r_{\perp}}(k=5)$ | $\sigma_{r_{\perp}}(k=50)$ | $\sigma_{r_{\perp}}(k=200)$ | Shape $\rho$ |
| --- | --- | --- | --- | --- | --- |
| 1 | TP53 pathway | 0.0245 | 0.0168 | 0.0079 | 0.9986 |
| 1 | RAS/MAPK | 0.0190 | 0.0114 | 0.0037 | 0.9992 |
| 1 | PI3K/AKT/mTOR | 0.0166 | 0.0118 | 0.0052 | 0.9995 |
| 2 | Immune checkpoint | 0.0033 | 0.0011 | 0.0003 | 1.0000 |
| 2 | BRCA DNA repair | 0.0182 | 0.0037 | 0.0014 | 0.9994 |
| 2 | Cell cycle/CDK | 0.0184 | 0.0035 | 0.0013 | 0.9992 |
| 3 | EMT | 0.0134 | 0.0039 | 0.0018 | 0.9998 |
| 3 | Angiogenesis | 0.0157 | 0.0030 | 0.0011 | 0.9996 |
| 3 | Apoptosis/BCL2 | 0.0200 | 0.0104 | 0.0047 | 0.9990 |
| 4 | Warburg effect | 0.0257 | 0.0126 | 0.0043 | 0.9986 |
| 4 | Stemness | 0.0123 | 0.0071 | 0.0032 | 0.9996 |
| 4 | NF-κB | 0.0135 | 0.0054 | 0.0016 | 0.9988 |
| 5 | Fenton metabolic | 0.0165 | 0.0069 | 0.0024 | 0.9992 |
| 5 | ceRNA hubs | 0.0228 | 0.0149 | 0.0065 | 0.9988 |
| 5 | Autophagy | 0.0205 | 0.0204 | 0.0090 | 0.9986 |
| 5 | Ferroptosis | 0.0240 | 0.0165 | 0.0083 | 0.9982 |
| C | Housekeeping | 0.0243 | 0.0095 | 0.0023 | 0.9989 |
| Summary across 17 entries | | $\sigma_{50}$ median 0.0104 | | | median 0.9992 |

**Table S15:**
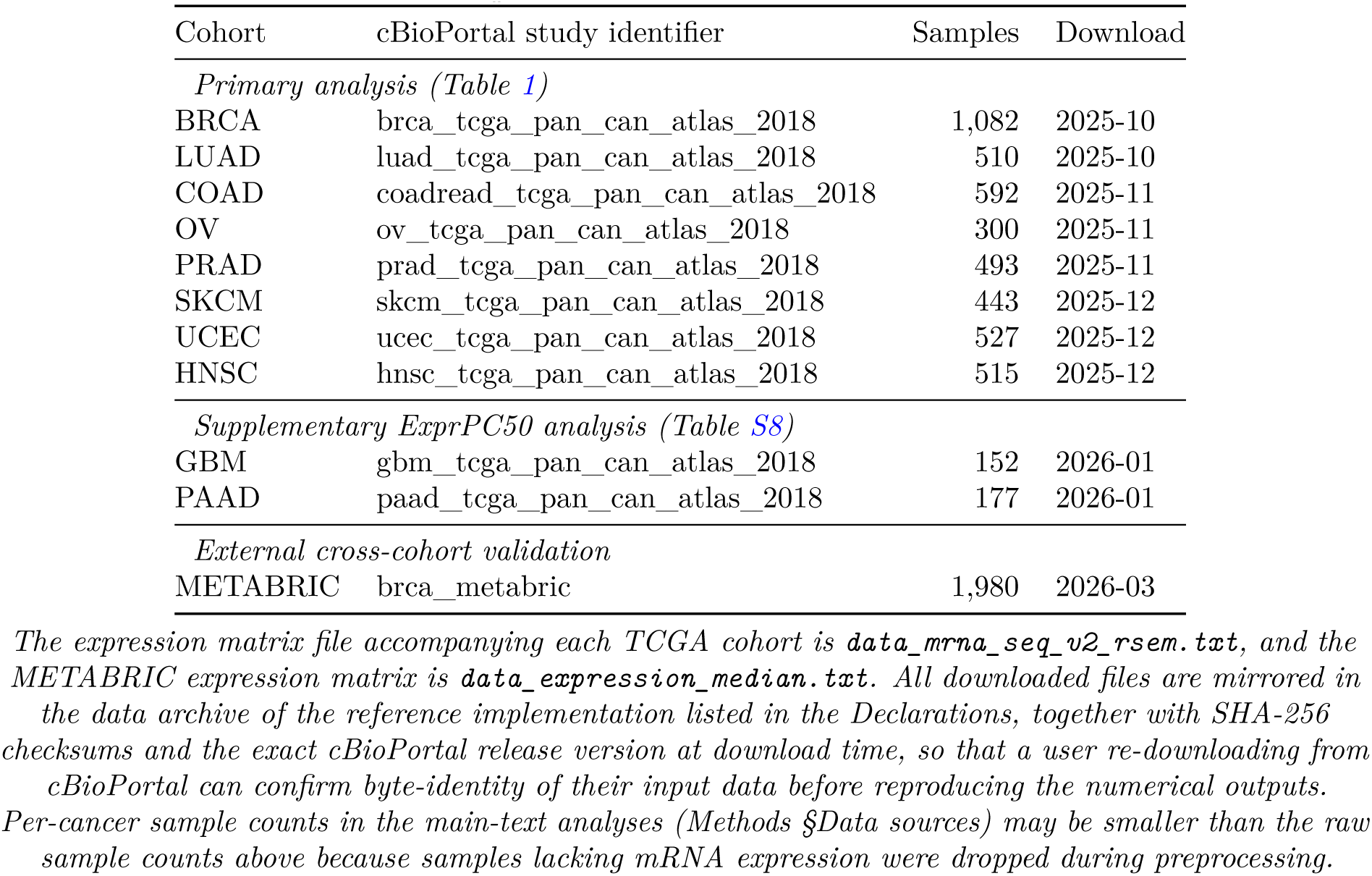
Cancer cohort study identifiers, release dates, and file provenance used in this work. All cohorts were downloaded from cBioPortal between October 2025 and March 2026 using the PanCancer Atlas 2018 data release for the ten TCGA studies and the METABRIC cohort for external validation. The eight primary-analysis cancer types in the upper block were used for the main Table 1 and all pan-cancer figures; the two supplementary cancer types (GBM, PAAD) in the middle block were used only for the ExprPC50 supplementary analysis in Supplementary Table S8; the METABRIC cohort in the bottom block was used only for the external cross-cohort validation reported in the main text.

| Cohort | cBioPortal study identifier | Samples | Download |
| --- | --- | --- | --- |
| *Primary analysis (Table 1)* |  |  |  |
| BRCA | brca_tcga_pan_can_atlas_2018 | 1,082 | 2025-10 |
| LUAD | luad_tcga_pan_can_atlas_2018 | 510 | 2025-10 |
| COAD | coadread_tcga_pan_can_atlas_2018 | 592 | 2025-11 |
| OV | ov_tcga_pan_can_atlas_2018 | 300 | 2025-11 |
| PRAD | prad_tcga_pan_can_atlas_2018 | 493 | 2025-11 |
| SKCM | skcm_tcga_pan_can_atlas_2018 | 443 | 2025-12 |
| UCEC | ucec_tcga_pan_can_atlas_2018 | 527 | 2025-12 |
| HNSC | hnsc_tcga_pan_can_atlas_2018 | 515 | 2025-12 |
| *Supplementary ExprPC50 analysis (Table S8)* |  |  |  |
| GBM | gbm_tcga_pan_can_atlas_2018 | 152 | 2026-01 |
| PAAD | paad_tcga_pan_can_atlas_2018 | 177 | 2026-01 |
| *External cross-cohort validation* |  |  |  |
| METABRIC | brca_metabric | 1,980 | 2026-03 |

The expression matrix file accompanying each TCGA cohort is `data_mrna_seq_v2_rsem.txt`, and the METABRIC expression matrix is `data_expression_median.txt`. All downloaded files are mirrored in the data archive of the reference implementation listed in the Declarations, together with SHA-256 checksums and the exact cBioPortal release version at download time, so that a user re-downloading from cBioPortal can confirm byte-identity of their input data before reproducing the numerical outputs. Per-cancer sample counts in the main-text analyses (Methods §Data sources) may be smaller than the raw sample counts above because samples lacking mRNA expression were dropped during preprocessing.

**Table S16:**
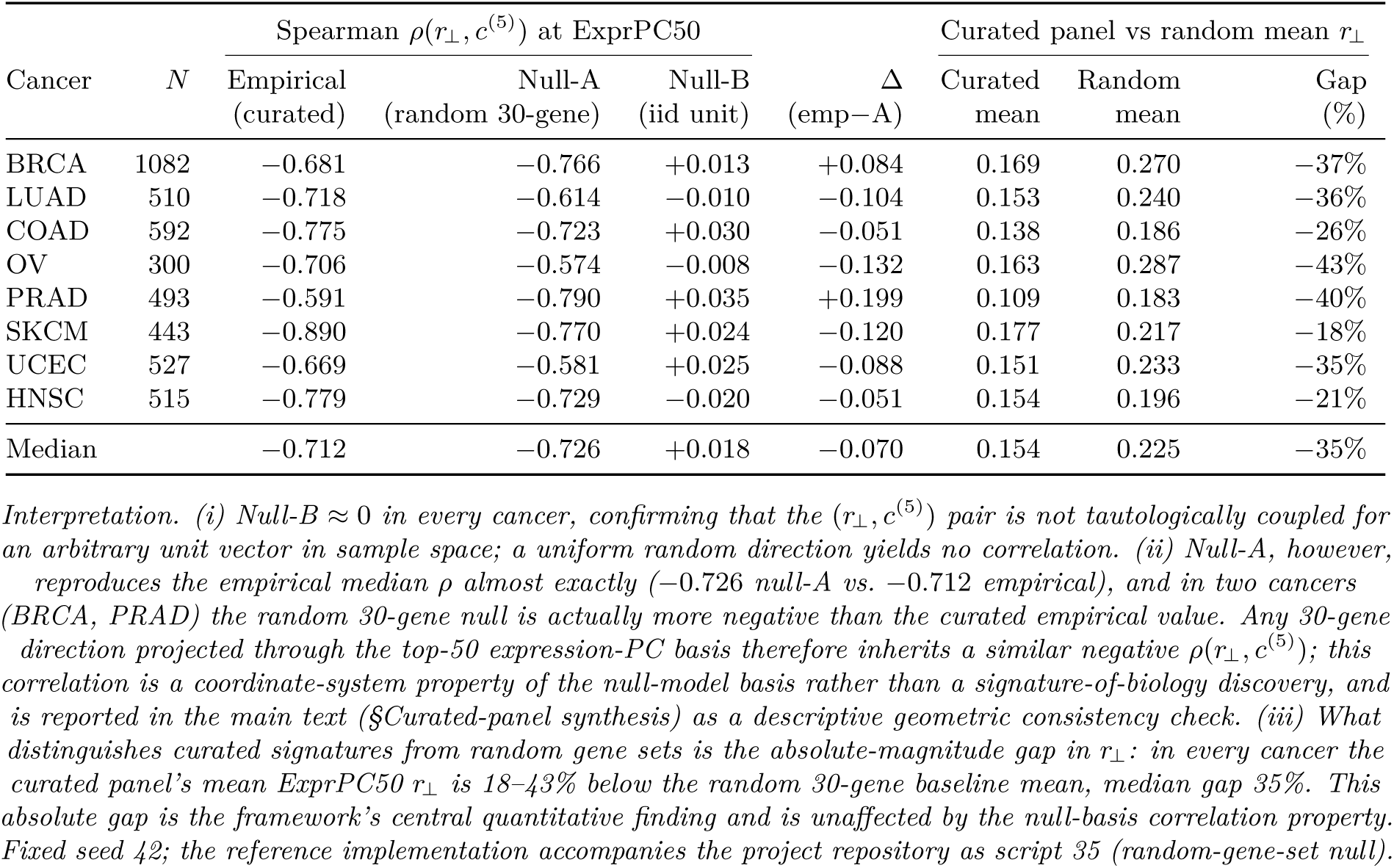
Random-gene-set null for the ExprPC50 *r*_⊥_-vs-*c*^(5)^ correlation, and curated-panel absolute gap vs random baselines. For each of the 8 primary-analysis TCGA cancer cohorts, we drew *B* = 1,000 random 30-gene sets from the gene universe of the preprocessed expression matrix and computed, for each draw, the residual ratio *r*_⊥_(*k* = 50) and the top-5 absorption concentration *c*^(5)^ under the same top-50 sample-space PC basis used for the curated benchmark (Methods §Statistical analysis; reference implementation accompanies the project repository as script 35). Column “Empirical curated *ρ*” repeats the 17-signature Spearman *ρ* between *r*_⊥_ and *c*^(5)^ reported in the main text and in Supplementary Table S10; column “Null-A *ρ* (random 30-gene)” gives the Spearman *ρ* across the 1,000 random-draw (*r*_⊥_*, c*^(5)^) pairs per cancer; column “Δ (emp null-A)” reports the difference. For reference, column “Null-B *ρ*” gives the corresponding Spearman *ρ* across 1,000 iid Gaussian unit vectors **h** (0*, I_N_*) in sample space (a uniform random direction that does not inherit the expression covariance geometry). Rightmost columns compare the curated 17-entry panel’s mean *r*_⊥_ at ExprPC50 to the random 30-gene baseline mean, both in absolute units and as a percentage gap; this magnitude gap is the quantitative discrimination between curated biological signatures and arbitrary gene combinations on which the framework’s central claim rests.

**Table S17:** Purity-aware null for the curated-panel ExprPC50 *r*_⊥_-vs-*c*^(5)^ correlation. For each cancer we rebuild a rank-50 sample-space basis as follows. Columns 1–3 are the PC1 directions of: an immune-infiltrate proxy panel (CD3D, CD3E, CD4, CD8A, CD8B, CD19, CD68, PTPRC, FOXP3, IFNG); a stromal/fibroblast proxy panel (COL1A1, COL1A2, COL3A1, VIM, FN1, ACTA2, PDGFRA, PDGFRB); and the 50 proliferation markers already used in the null-model hierarchy (§Null model hierarchy). Columns 4–50 are the top-47 PCs of the residual expression matrix *Y* − *Q*_bio_*Q*^𝖳^ *Y* after QR-orthonormalization of the 3-column biological block; the full 50-column basis is re-orthonormalized by a final QR pass. The 17 curated benchmark entries are then re-scored under this purity-aware basis, and the per-cancer Spearman *ρ* between *r*_⊥_ and *c*^(5)^ is recomputed. Column “Standard *ρ*” is the value reported in the main text (same as Supplementary Table S16, column “Empirical curated”); column “Purity-adjusted *ρ*” is the value under the new basis; column “Δ” is the difference. This test was pre-specified as a check on whether the observed *ρ*(*r*_⊥_*, c*^(5)^) could be driven by tumor-purity, immune, or stromal composition artifacts that a standard top-50 PC basis would implicitly absorb.

| Cancer | $N$ | Standard $\rho$ | Purity-adj $\rho$ | $\Delta$ | Bio-block columns kept |
| --- | --- | --- | --- | --- | --- |
| BRCA | 1082 | −0.681 | −0.907 | −0.226 | immune, stromal, prolifer |
| LUAD | 510 | −0.718 | −0.890 | −0.172 | immune, stromal, prolifer |
| COAD | 592 | −0.775 | −0.860 | −0.086 | immune, stromal, prolifer |
| OV | 300 | −0.706 | −0.775 | −0.069 | immune, stromal, prolifer |
| PRAD | 493 | −0.591 | −0.865 | −0.275 | immune, stromal, prolifer |
| SKCM | 443 | −0.890 | −0.853 | +0.037 | immune, stromal, prolifer |
| UCEC | 527 | −0.669 | −0.757 | −0.088 | immune, stromal, prolifer |
| HNSC | 515 | −0.779 | −0.801 | −0.022 | immune, stromal, prolifer |
| Median |  | −0.712 | −0.857 | −0.087 | — |

**Table S18:** Partial Spearman (size-controlled) and Shannon-entropy companion diagnostics. Left block: partial Spearman *ρ*(*r*_⊥_*, c*^(5)^ *n*_genes_) controlling for gene-set size, alongside the marginal Spearman reported in the main text (Supplementary Table S10). Partial correlation is computed by regressing ranks of *r*_⊥_ and *c*^(5)^ each on ranks of *n*_genes_ linearly and Pearson-correlating the residuals (standard partial-Spearman definition). Right block: Shannon entropy *H*(**a**) =Σ*_j_ p_j_* log *p_j_* (nats) with *p_j_* = *a_j_/*Σ *i a_i_*across the top-50 PC absorption spectrum, reported as an information-theoretic companion to the inverse participation ratio (IPR). Columns “*ρ*(*r*_⊥_*, H*)” and “*ρ*(*H,* IPR)” give the per-cancer Spearman correlation between *r*_⊥_ and Shannon entropy (should be strongly positive if the Shannon statistic agrees with IPR on the same phenomenon) and between Shannon entropy and IPR (should be ≈ 1 if the two diagnostics are rank-equivalent on this panel).

| Cancer | $N$ | Size-controlled Spearman | | | Shannon entropy | |
| --- | --- | --- | --- | --- | --- | --- |
| | | Marginal $\rho$<br>($r_{\perp}, c^{(5)}$) | Partial $\rho$<br>($\vert n_{\text{genes}}$) | $\Delta$<br>(part – marg) | $\rho(r_{\perp}, H)$ | $\rho(H, \text{IPR})$ |
| BRCA | 1082 | −0.681 | −0.700 | −0.019 | +0.645 | +0.900 |
| LUAD | 510 | −0.718 | −0.718 | +0.000 | +0.657 | +0.944 |
| COAD | 592 | −0.775 | −0.813 | −0.038 | +0.750 | +0.990 |
| OV | 300 | −0.706 | −0.730 | −0.024 | +0.650 | +0.951 |
| PRAD | 493 | −0.591 | −0.650 | −0.059 | +0.588 | +0.922 |
| SKCM | 443 | −0.890 | −0.890 | +0.000 | +0.907 | +0.961 |
| UCEC | 527 | −0.669 | −0.773 | −0.104 | +0.696 | +0.924 |
| HNSC | 515 | −0.779 | −0.782 | −0.003 | +0.792 | +0.993 |
| Median |  | −0.712 | −0.752 | −0.022 | +0.676 | +0.947 |

**Table S19:** Biological annotation of the top-10 expression PCs in each TCGA cohort. For each of the 8 primary-analysis cancers, we correlate the leading left singular vectors *U*:,_1_, …, *U*_:,10_ of the centered expression matrix (the same PC scores used to construct the null-model basis) against three per-sample biological proxy scores: mean expression across a 10-gene immune-infiltrate panel (CD3D, CD3E, CD4, CD8A, CD8B, CD19, CD68, PTPRC, FOXP3, IFNG), mean expression across an 8-gene stromal/fibroblast panel (COL1A1, COL1A2, COL3A1, VIM, FN1, ACTA2, PDGFRA, PDGFRB), and mean expression across the 50 proliferation markers already used in the null-model hierarchy. Columns “Top-PC” report the PC index most strongly correlated with each proxy (by *ρ*) and the signed Spearman correlation at that index; column “%_PC1_” reports the fraction of total variance explained by the first PC.

| Cancer | $N$ | %PC1 | Top-PC immune | Top-PC stromal | Top-PC proliferation |
| --- | --- | --- | --- | --- | --- |
| BRCA | 1082 | 12.2% | PC1 (+0.58) | PC2 (+0.65) | PC2 (−0.52) |
| LUAD | 510 | 9.9% | PC2 (−0.56) | PC2 (−0.62) | PC1 (−0.77) |
| COAD | 592 | 13.2% | PC1 (−0.67) | PC1 (−0.86) | PC2 (+0.34) |
| OV | 300 | 9.5% | PC1 (−0.72) | PC2 (+0.69) | PC3 (+0.58) |
| PRAD | 493 | 15.2% | PC1 (−0.64) | PC1 (−0.67) | PC3 (−0.50) |
| SKCM | 443 | 11.3% | PC1 (+0.70) | PC3 (−0.43) | PC7 (+0.48) |
| UCEC | 527 | 9.2% | PC4 (+0.66) | PC4 (+0.53) | PC1 (+0.64) |
| HNSC | 515 | 12.2% | PC4 (+0.80) | PC2 (−0.65) | PC5 (−0.42) |

